# FAM136A is an essential chaperone for mitochondrial membrane protein biogenesis

**DOI:** 10.64898/2026.08.03.742567

**Authors:** Melanie Ernst, Jichen Zhang, Haoqi Xu, Alexa Ma, Lena A. K. Bögeholz, Marton Szabo, Ting-Yu Wang, Tsui-Fen Chou, Alina Guna, Rebecca M. Voorhees

## Abstract

The metabolic and signaling function of mitochondria rely on a network of chaperones within the inner membrane space (IMS) that regulate the biogenesis of nascent mitochondrial proteins. Using a genome wide CRISPRi screen we found that in human cells FAM136A is required for biogenesis of all three voltage-dependent anion channel (VDAC) paralogs, an abundant and essential family of β-barrel metabolite transporters in the outer mitochondrial membrane (OM). FAM136A is a ubiquitously expressed essential gene, that is conserved in metazoa and plants. Using a combination of experiments in human cells and in vitro reconstitution, we determined that FAM136A associates with unfolded VDACs in the IMS; solubilizes nascent VDAC through a direct interaction; and facilitates insertion of VDAC into the OM. FAM136A also binds and chaperones a subset of α-helical subunits of the electron transport chain. We therefore conclude that FAM136A is an IMS-resident chaperone, necessary and sufficient to maintain nascent membrane proteins in a folding-competent state to mediate their integration into the bilayer.

## INTRODUCTION

Despite two billion years of co-evolution^1^, the endosymbiotic origin of mitochondria continues to define the biogenesis of many mitochondrial proteins. Though many of the core pathways are conserved, comparative genomics suggests that the mammalian mitochondrial proteome is over 60% larger than in fungi ^2,3^, due primarily to protein expansion in metazoans and ancestral protein loss in fungus^4^. This increased complexity creates a commensurate burden on the biogenesis machinery, which must simultaneously accommodate a larger number of substrates and more biophysical and topological diversity. How mammalian mitochondria have evolved to handle this proteome expansion remains incompletely understood.

The impact of mitochondria’s evolution on protein biogenesis is particularly evident for β -barrel proteins, which are found only in the outer mitochondrial membrane (OM), and play critical roles in metabolite transport, protein import, and membrane protein biogenesis. Although all mitochondrial outer membrane (OM) proteins are now encoded in the nuclear genome^5^, integration of β-barrel proteins still relies on machinery inherited from their ancestral bacterial counterparts.

In gram-negative bacteria, OM β-barrels are synthesized in the cytosol, translocated across the inner membrane, and ferried through the periplasmic space by a suite of soluble chaperones^6^. Subsequent delivery to the OM and the β-barrel assembly machinery (BAM) complex, which contains a central β-barrel core subunit, BamA, results in templated insertion and folding of the nascent β-barrel into the OM^7–9^. Similarly, in mitochondria, β-barrels are first translated in the cytosol, and then translocated across the OM by the translocase of the outer membrane (TOM) complex into the intermembrane space (IMS), the equivalent of the ancestral periplasm^10–13^. Here, in the aqueous environment of the IMS, β-barrels pose a major challenge, as they contain β-strands with alternating hydrophobic (lipid-facing in the final fold) and hydrophilic or small polar residues (protein-facing in the final fold)^14,15^. These hydrophobic residues make β-barrels prone to aggregation in the aqueous environment of the IMS^16,17^. As they transit through the IMS, the nascent, unfolded proteins must therefore be chaperoned similar to the β-barrels in the periplasm, preventing aggregation and remaining import competent by avoiding non-productive interactions^18^. Experiments primarily in yeast mitochondria have shown that the small translocase of the inner membrane (Tim) proteins (Tim9-10 and Tim8-13) function as soluble chaperones in the IMS^10,19^. In the aqueous environment of the IMS the Tims bind to nascent β-barrel substrates to shield their hydrophobic stretches and prevent aggregation. The soluble Tims have also been shown to bind and chaperone inner membrane α-helical proteins ^20–23^. At the OM, nascent β-barrel substrates are delivered to the eukaryotic homolog of BAM, the sorting and assembly machinery (SAM) complex, for templated integration into the OM^24,25^. The extent to which these observations apply to mammalian systems is not clear.

Emblematic of the proteome differences between fungus and mammals are the voltage dependent anion channels (VDACs), which are the most abundant proteins of the OM ^26,27^. VDACs adopt a β-barrel topology composed of 19 strands, which provides the primary pathway for transport of small molecule metabolites between the cytosol and mitochondria. While fungi contain one to two such channels, Porin1 and Porin2, metazoans require three to four separate paralogs while plants express up to ten such channels^28^. Despite similarity in their overall structure, the three human paralogs VDAC1,2, and 3 have distinct sequences and properties, differential tissue expression, and play unique biological roles in both normal and pathological conditions^29^. While all three paralogs can conduct metabolites and ions, VDAC1 specifically plays a role in apoptotic calcium signaling. In contrast, VDAC3 protects cells from accumulation of reactive oxygen species (ROS) and regulates the electron transport chain (ETC) and thereby respiration, particularly in cellular states with high energy demands ^29^. This increased substrate diversity may place additional demands on the mammalian β-barrel biogenesis machinery. To accommodate these additional substrates, we hypothesized that mammalian mitochondria may possess specialized biogenesis factors which we set out to identify and characterize.

### Systematic identification of factors required for β-barrel biogenesis in human mitochondria

To study β-barrel biogenesis in human cells, we chose to initially focus on the VDACs due to their expansion in metazoa, their essential function, and their high expression relative to other OM proteins. To develop a fluorescent reporter system to monitor nascent VDAC expression, we leveraged a split GFP approach composed of three components: an IMS-localized GFP1-10 composed of the first ten β-strands of GFP; a GFP11-fusion of the 11^th^ β-strand of GFP to the N-termini of VDAC1,2, and 3; and finally, an RFP normalization control (**Fig. 1A**)^30,31^. Using this strategy, we verified that all three GFP-11-tagged VDAC constructs were integrated into mitochondria in a TOM and SAM dependent manner, resulting in GFP complementation and fluorescence (**Extended Data Fig. 1A and C**). Surprisingly however, when we tested the effects of depletion of each of the IMS-localized small TIM proteins on these reporters, we observed only a modest effect on any VDAC paralog (**Fig. 1B**, **Extended Data Fig. 1B and C**). No statistically significant effect of depletion of any TIM protein was observed for mitochondrial integration of VDAC3 (**Fig. 1B**). Given its unique role in electron transport chain (ETC) maintenance and apparent dependence on an unidentified IMS chaperone for biogenesis, we therefore selected VDAC3 for further analysis.

**Figure 1.**
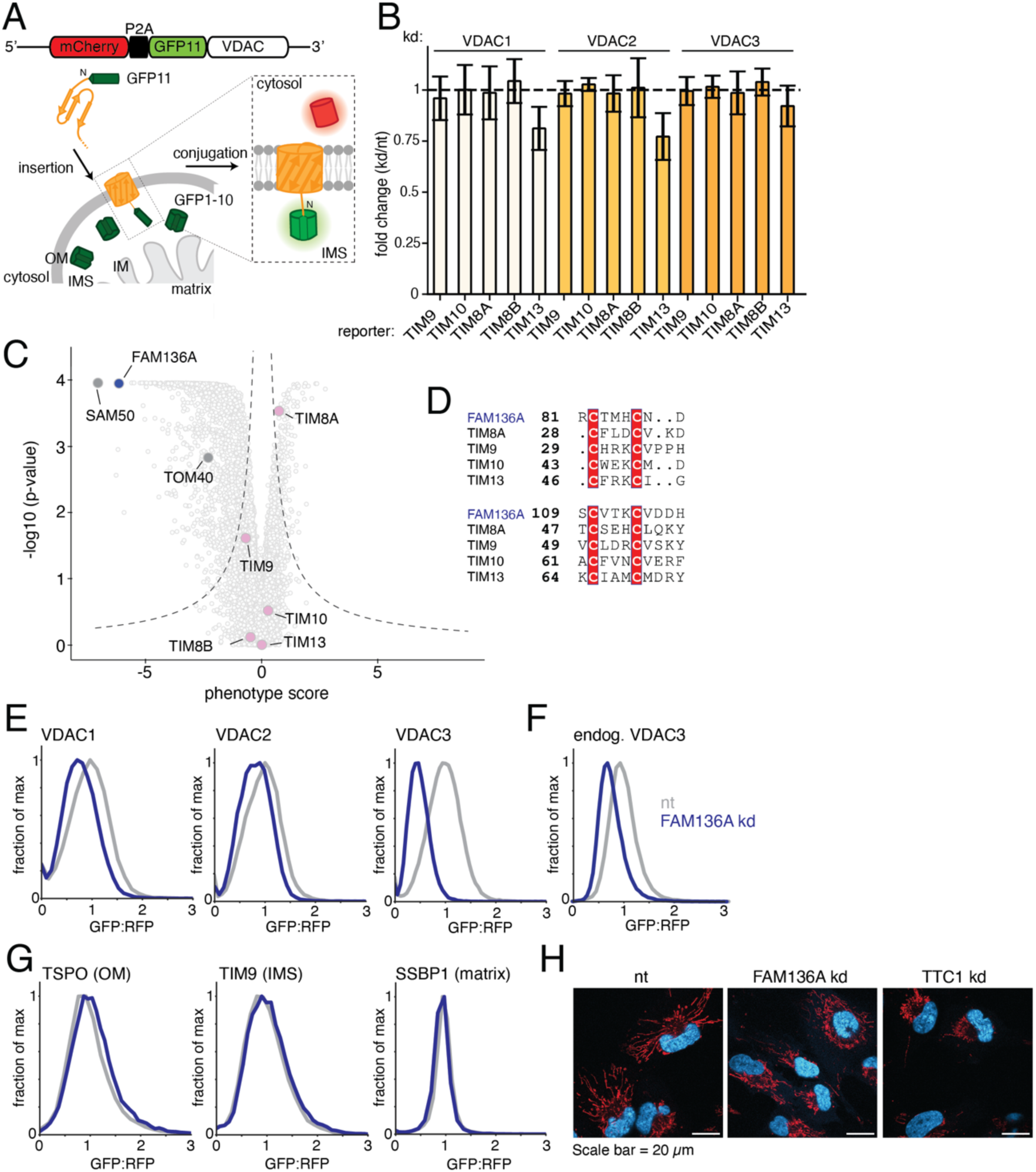
FAM136A is required for VDAC biogenesis in human cells. **(A)** Schematic of the split GFP reporter system used to probe the biogenesis and localization of nascent mitochondrial proteins to the mitochondria^30,31^. The first ten β-strands of GFP (GFP1-10) were constitutively targeted to the intermembrane space (IMS), while the 11^th^ β-strand (GFP11) was fused to the appropriate termini of a mitochondrial substrate, such as the N-terminus of VDAC3. Successful mitochondrial integration, exposing GFP11 to the IMS, resulted in GFP complementation and thereby fluorescence. Expression of this GFP11-tagged reporter from a single open reading frame along with a normalization marker (RFP) specifically allows the query of post-translational processes. Note, using this reporter system alone, for outer membrane (OM) substrates such as the VDACs, we cannot distinguish between nascent proteins that have been translocated into the IMS from those that are integrated into the OM. However, experiments using the three mammalian VDAC paralogs (VDAC1-3) indicated that depletion of factors that trap nascent substrates in the cytosol (e.g. TOM40) or in the IMS (e.g. SAM50), both result in a decrease in GFP:RFP fluorescence (Extended Data Fig. 1A-B). We therefore concluded that our platform faithfully reported on factors required throughout the maturation of VDACs. **(B)** The reporter system described in (A) was used to test the dependence of the three mammalian VDAC paralogs on the known IMS-resident chaperones, translocase of inner membrane (TIM) TIM9/10 and TIM8/13. Mitochondrial integration of GFP11-VDAC1,2, and 3 was assessed in human K562 CRISPRi cells upon depletion of TIM8A, TIM8B, TIM9, TIM10, TIM13 (kd) compared to a non-targeting control (nt). Shown is the fold change for each VDAC reporter calculated as (GFP:RFP in kd cells)/(GFP:RFP in nt cells). A fold change of 1 indicates no change compared to the nt control. Shown is the mean of 2-3 independent biological replicates ± SD. Western blots assessing TIM depletion are shown in Extended Data Fig. 1C. **(C)** Systematic analysis of VDAC3 biogenesis in human cells was performed using a genome wide CRISPRi screen using the VDAC3 reporter depicted in (A). K562 CRISPRi cells stably expressing GFP11-VDAC3 along with the normalization marker (RFP) were transduced with the TOP5 v2 sgRNA library. Following eight days of depletion, cells with altered GFP:RFP ratios were isolated using FACS and deep sequenced to identify enriched sgRNAs in the top (increased GFP:RFP) and bottom (decreased GFP:RFP) populations. Depicted is a volcano plot of the phenotype score [log2(increased GFP:RFP/decreased GFP:RFP)] for the three strongest sgRNAs versus the -log of the Mann-Whitney p values. Genes that fall outside the dashed lines represent statistically significant hits. Individual genes are displayed in light grey, the biogenesis machineries TOM40 and SAM50 are highlighted in dark grey, the IMS-resident TIM chaperones in pink, and FAM136A is shown in blue. **(D)** Sequence alignment comparing the twin CX_3_C motifs from human FAM136A, TIM9, TIM10, TIM8A, and TIM13. The two characteristic cysteines of the motif are highlighted in red. The numbers on the left indicate the amino acid position. **(E)** Integration into mitochondria of the GFP11-VDAC reporters described in (A) was assessed in K562 CRISPRi cells expressing GFP1-10 in the IMS and a non-targeting control (nt) or FAM136A knock down sgRNA (kd). GFP fluorescence relative to a normalization marker was determined by flow cytometry and displayed as a histogram. Western blots assessing FAM136A depletion are shown in Extended Data Fig. 1C. **(F)** To exclude potential artifacts from overexpression, we introduced an RFP_P2A_GFP11 tag at the N-terminus of VDAC3 at its endogenous locus in K562 CRISPRi cells (described in detail in the materials and methods). Mitochondrial integration of endogenous levels of VDAC3 was measured relative to an expression control (RFP) in FAM136A depleted (kd) compared to control (nt) cells. GFP fluorescence relative to a normalization marker was determined by flow cytometry and displayed as a histogram. **(G)** To test whether defects in VDAC biogenesis upon loss of FAM136A were specific, we generated reporters for several representative mitochondrial proteins. Integration of TSPO (an α-helical OM protein), TIM9 (an IMS-resident soluble protein), and SSBP1 (a soluble matrix protein) were assessed as described in (E). **(H)** Confocal microscopy of mitochondrial morphology in RPE1 cells transduced with either a non-targeting sgRNA (nt), FAM136A sgRNA (FAM136A kd) or the positive control TTC1 sgRNA (TTC1 kd)^64^. Staining is shown in for DAPI (blue; nucleus) and MitoTracker (red; mitochondria). For comparison, mitochondrial fragmentation in cells transduced with a TTC1 sgRNA is shown. Mitochondrial fragmentation is characterized by loss of extended red-stained mitochondrial structures.

Using our validated ratiometric VDAC3 reporter, we performed a genome-wide CRISPR interference (CRISPRi) screen using the Top5 v2 sgRNA library^32,33^. Consistent with our earlier analysis, while depletion of SAM50 and TOM40 had pronounced effects, none of the TIM proteins were implicated as biogenesis factors in our screen. (**Fig. 1C**). However, analysis of the statistically significant hits identified a protein of unknown function, FAM136A, that contains two twin-CX_3_C motifs, which are also found in the small TIM proteins and other MIA40-dependent substrates (**Fig. 1D, Extended Data Fig. 2A**)^34–36^. FAM136A is a common essential gene^37,38^, previously shown to be localized to the IMS, a dimer, a MIA40 substrate (**Extended Data Fig. 2B-D**) and broadly expressed across tissue types^34,35^. It is conserved across some holozoa and plants, but is not found in fungi, suggesting it may have been evolutionarily lost in the fungal lineage (**Extended Data Fig. 3**). Earlier experiments found that depletion of FAM136A results in induction of the integrated stress response and decreases mitochondrial ATP production, underscoring its critical role in mitochondrial proteostasis^35^. In patients, a premature stop codon in FAM136A causes the autosomal-dominant Ménière’s disease, characterized by sensorineural hearing loss^39^. However, its molecular function, and therefore a biochemical explanation for these phenotypes was not known.

In a subsequent arrayed reporter screen, we found that depletion of FAM136A most prominently affected VDAC3, though it decreased biogenesis of all three VDAC paralogs (**Fig. 1E and Extended Data Fig. 1C**). To exclude artifacts from overexpression, we verified that the levels of endogenously expressed VDAC3 were also decreased by loss of FAM136A (**Fig. 1F**). In contrast, its depletion did not affect several IMS, OM, and matrix localized controls (**Fig. 1G)** or visibly altered mitochondrial morphology (**Fig. 1H**). Together, these experiments excluded a generalized defect in mitochondrial function in the absence of FAM136A, pointing to a more specific role in membrane protein biogenesis.

### FAM136A is an IMS-localized chaperone for VDAC biogenesis

Given its localization to the IMS and the presence of a twin-CX_3_C motif, we hypothesized that FAM136A could function as an IMS chaperone for VDAC biogenesis. We reasoned that for this to be true, FAM136A must have the following features: (i) it physically associates with unfolded VDAC in the IMS, (ii) it is sufficient to solubilize nascent VDAC through a direct interaction, and (iii) it facilitates insertion into the OM. We set out to systematically test these features using a combination of experiments in human cells and in vitro reconstitution.

#### FAM136A physically associates with the VDACs in the IMS

To determine if FAM136A can capture the VDAC paralogs, we first tested whether it binds endogenous VDACs in cells. For this, we generated a stable cell line that exogenously expressed ALFA-tagged FAM136A, which remains functional in VDAC biogenesis (**Extended Data Fig. 4A**). Immunoprecipitation of FAM136A under native conditions from mitochondria revealed that all three endogenous VDACs are among the most enriched hits by mass spectrometry (**Fig. 2A**). Accordingly, three bands consistent in size with the three VDAC paralogs could be directly visualized in the FAM136A immunoprecipitation by SDS-PAGE, which we confirmed corresponded to VDAC1,2, and 3 by western blot (**Fig. 2B**). We concluded that this interaction was specific because matched immunoprecipitations of either TIM9 or TIM10 did not appreciably co-purify any of the VDAC paralogs (**Fig. 2B**).

**Figure 2:**
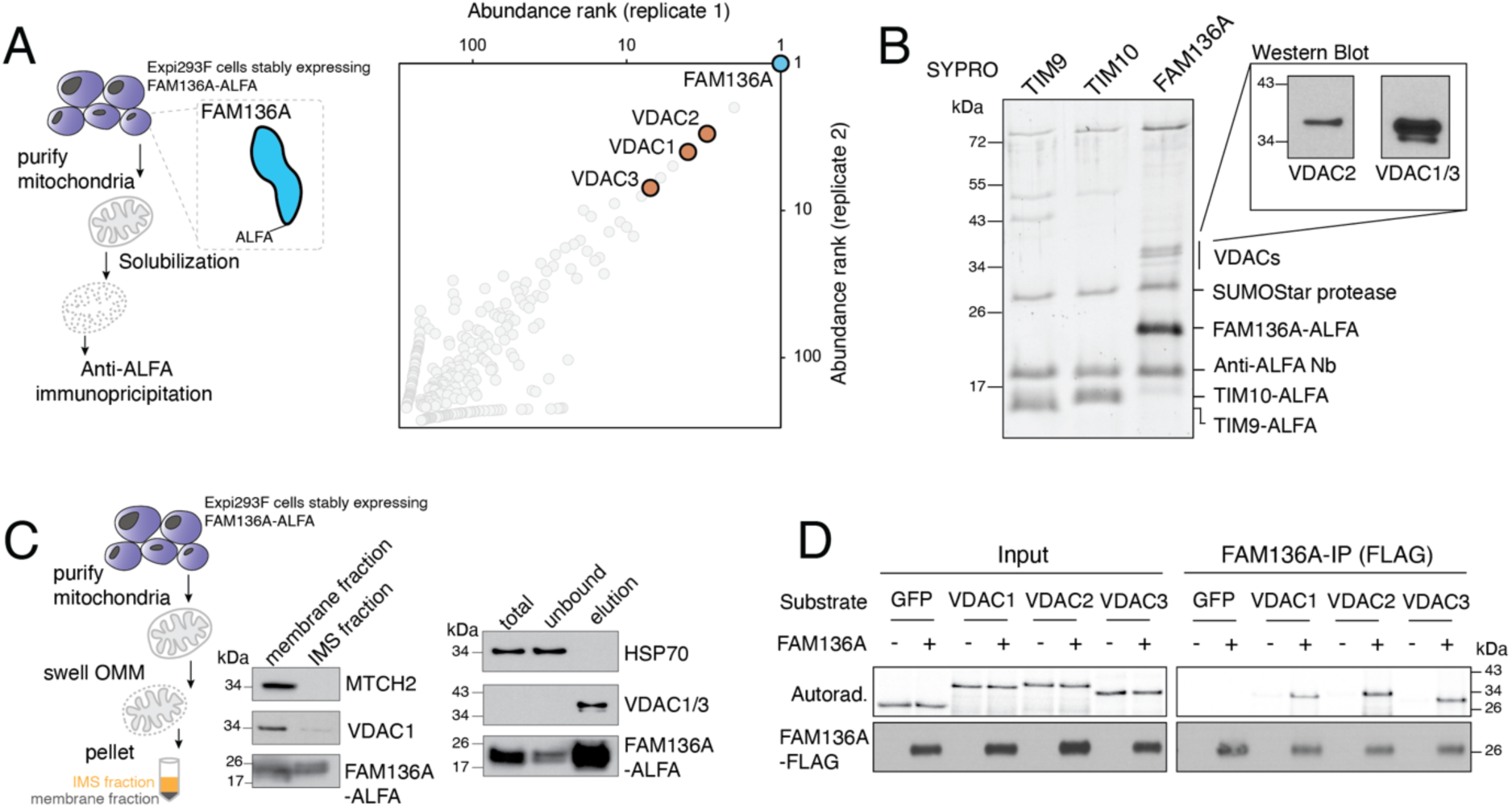
FAM136A interacts with nascent VDACs in the IMS. **(A)** Analysis of the interactome of exogenously expressed FAM136A using mass spectrometry. FAM136A was isolated under native conditions following solubilization in 1% GDN from the mitochondrial fraction of an Expi293F cell line stably expressing FAM136A-ALFA. Co-purifying proteins were ranked by abundance (intensity-based absolute quantification, iBAQ), with one of two biological replicates shown on each axis. Co-purification of the three endogenous VDAC isoforms are highlighted in orange, and FAM136A-ALFA is in blue. **(B)** Exogenously expressed TIM9-ALFA, TIM10-ALFA, and FAM136A-ALFA were purified from Expi293F cells following solubilization in 1% GDN and eluted by on-column protease cleavage. Co-purifying proteins were analyzed by SDS-PAGE and stained by Sypro Ruby. Tagged proteins, anti-ALFA nanobody, and the SUMOStar protease are annotated. The FAM136A immunoprecipitations were further analyzed by western blot using antibodies against VDAC2 and VDAC1/3. **(C)** To test if FAM136A could capture the population of unfolded VDAC within the IMS, we performed an immunoprecipitation of FAM136A as in (A) with the following modifications. Mitochondria were purified from Expi293F cells stably expressing FAM136A-ALFA and the IMS fraction was isolated by swelling and rupture of the OM using hypotonic conditions followed by ultracentrifugation. Western blots for a representative OM protein (MTCH2) were used to verify quantitative depletion of the OM membrane from the IMS fraction. Samples from the membrane and IMS fraction were also analyzed by western blotting using an antibody that recognizes both VDAC1 and VDAC3 (VDAC1/3). In this exposure, we detect only the more prominent VDAC1 but find that small levels of endogenous VDAC1 are present in the IMS at steady-state. FAM136A-ALFA was subsequently immunoprecipitated from this purified IMS lysate and analyzed by SDS-PAGE and western blot, confirming the specific co-purification of endogenous VDAC1 and VDAC3 with FAM136A from the IMS fraction. **(D)** Analysis of nascent β-barrel association with FAM136A in vitro in the absence of mitochondria. The indicated ^35^S-methionine labeled substrates were translated in rabbit reticulocyte lysate (RRL) in the absence or presence of recombinant FAM136A-3xFLAG purified from *E. coli*. FAM136A was immunoprecipitated using anti-FLAG resin and eluted with FLAG peptide. Co-purification of each substrate was analyzed by SDS-PAGE and autoradiography. Samples were also subjected to western blotting to ensure an equal amount of FAM136A-3xFLAG in the input and elution.

While these experiments established an interaction between FAM136A and endogenous VDACs, they could not differentiate between binding to the mature β-barrel in the OM vs the nascent folding intermediate. Therefore, we tested if FAM136A could specifically capture the population of endogenous VDAC present at steady-state levels in the IMS (**Fig. 2C**). To do this we used our FAM136A-ALFA expressing cell line and subjected it to hypotonic lysis and fractionation to purify the IMS, verifying that we had quantitatively depleted the OM (**Fig. 2C**). Immunoprecipitation of FAM136A from this lysate again identified endogenous VDACs as specific interactors of FAM136A, consistent with its binding to the unfolded nascent β-barrel in the IMS (**Fig. 2C**).

Additionally, we tested if this interaction between FAM136A and the VDACs could occur in the absence of mitochondria. To do this, we expressed and purified FLAG-tagged FAM136A in *E. coli*, to ensure that no other mitochondrial or human proteins would contaminate the preparation (**Extended Data Fig. 4B**). Using a cell free rabbit reticulocyte lysate (RRL) translation system, devoid of mitochondria or any source of membranes, we expressed untagged VDAC1,2, and 3 in the presence of our recombinant FLAG-tagged FAM136A. We found that in the absence of any additional mitochondrial factors, FAM136A was sufficient to bind and immunoprecipitate all three VDAC paralogs, but not the soluble β−barrel, GFP (**Fig. 2D**). This interaction appeared to be highly specific, as reticulocyte extract contains many other proteins and chaperones. Furthermore, we found that truncations of VDAC3 that cannot fold into a complete β-barrel were still competent for binding to FAM136A (**Extended Data Fig. 4C-D**), consistent with the model that FAM136A could capture nascent VDAC as it transits into the IMS, but before the complete protein has translocated across the OM. Together, we therefore conclude that FAM136A can interact with the population of unfolded, nascent VDAC in the IMS.

#### FAM136A is sufficient to solubilize nascent VDAC3 via a direct interaction

However, because these experiments were performed in cells and extracts, we next sought to test whether FAM136A was sufficient to capture the VDACs. We would expect a chaperone to not only directly bind its substrates, but also to keep them soluble in an aqueous environment such as the IMS. To test this, we leveraged our cell free RRL translation system using either full-length or truncated VDAC3, the substrate where we see the largest dependence on FAM136A in cells (**Fig. 1E**). Despite the presence of many cytosolic chaperones in the translation extract^40^, ten or more strands of nascent VDAC3 aggregates and pellets on a sucrose gradient (**Fig. 3A-B**). In contrast, full-length GFP, a soluble β-barrel that does not require chaperoning migrates in the early fractions of the gradient (**Extended Data Fig. 5A**). However, addition of purified recombinant FAM136A solubilizes both the truncated and full-length VDAC3, and we observe FAM136A co-migrating in the VDAC-containing fractions of the gradient (**Fig. 3B**).

**Figure 3:**
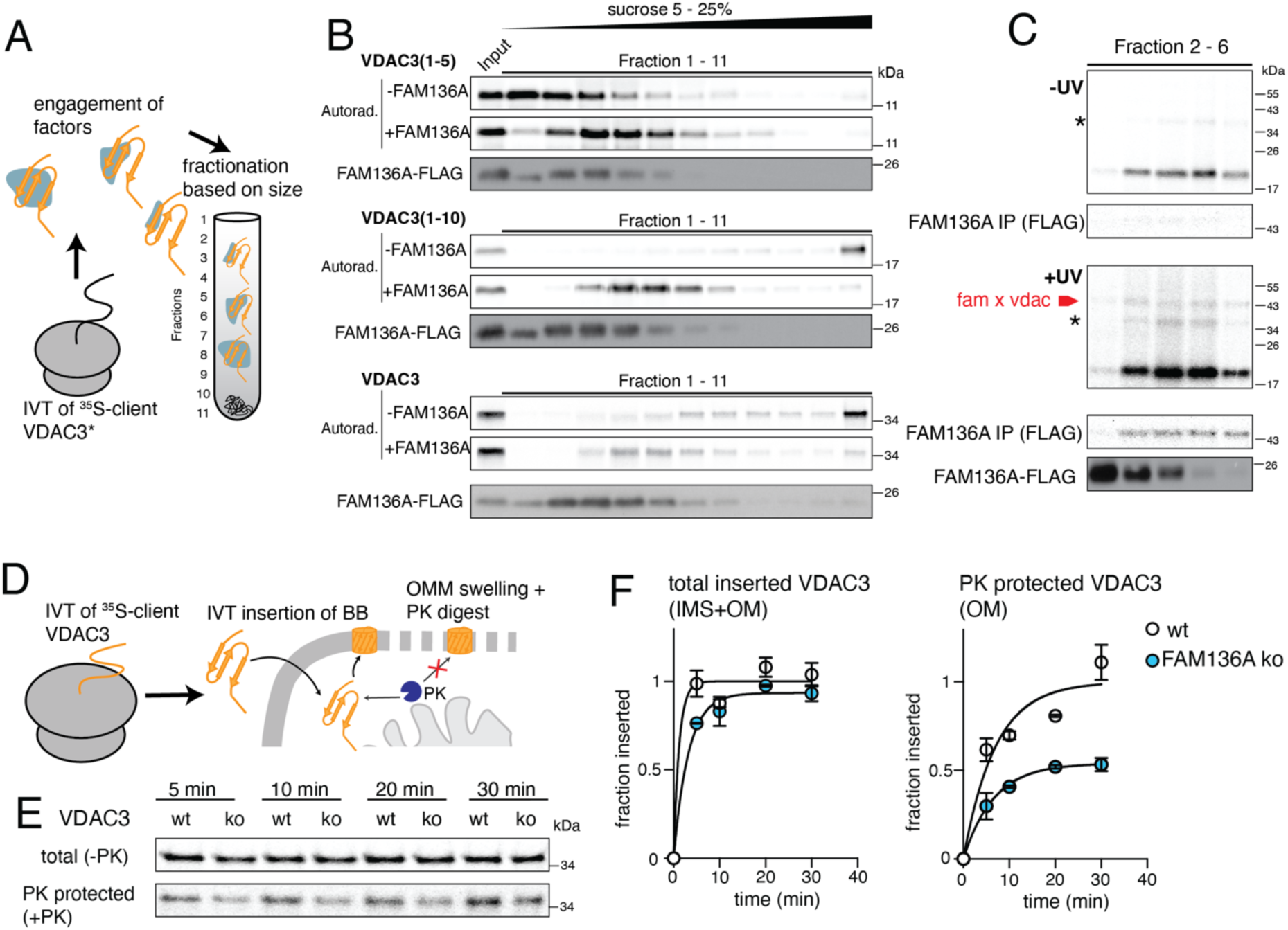
FAM136A is an IMS-localized chaperone that solubilizes VDAC3 and facilitates its insertion into the OM. **(A)** Schematic of the assay used to test if FAM136A is (i) sufficient to maintain nascent unfolded VDAC3 in a soluble form in an aqueous environment, and (ii) does so via a direct physical interaction. To site-specifically insert a crosslinker into nascent VDAC3, ^35^S-methionine labeled VDAC3 harboring an amber stop codon at I29 in the first β-sheet or at position N57 (in the third β-sheet) (VDAC3*) was translated in RRL supplemented with BpA-tRNA(UAG), the photoactivatable crosslinker BpA, and the BpA aminoacyl synthetase. The resulting VDAC3* containing the crosslinker BpA at a single site within the nascent β-barrel was translated in the absence or presence of recombinant FAM136A-3xFLAG. We reasoned that in the absence of mitochondria, unfolded VDAC would aggregate, but a direct interaction with an appropriate chaperone could maintain it in a soluble state. To test this hypothesis, we fractionated the resulting VDAC3* translation reaction over a sucrose gradient. We collected fractions from the top, where small soluble proteins would be expected to migrate (fraction 1), to the bottom, where insoluble proteins would be expected to pellet (fraction 11) ^65^. Each fraction was then exposed to UV light for crosslinking analysis and analyzed via SDS-PAGE and autoradiography. **(B)** Analysis of the solubility of VDAC3* in the absence or presence of FAM136A-3xFLAG. Full length VDAC3* (FL) and two truncated VDAC3* mutants (1-5, 1-10) were translated as ^35^S-methionine labeled substrates and fractionated on a sucrose gradient as described in (A). Nascent VDAC3 was visualized by autoradiography, while the migration of FAM136A-3xFLAG on the sucrose gradient was determined by western blotting. A control of FAM136A not solubilizing GFP is shown in Extended Data Fig. 5A. **(C)** Site-specific UV crosslinking of VDAC3* harboring an amber stop codon at I29 in the first β-sheet and FAM136A. Fractions of VDAC3* (1-10) translated with FAM136A-3xFLAG from (B) were subjected to photo crosslinking under UV light, followed by a FLAG-IP under denaturing conditions to enrich for FAM136A crosslinked species. Samples were analyzed by SDS-PAGE and autoradiography. Shown are fractions 2 to 6 and results for crosslinking at position N57 (in the third β-sheet) are shown in Extended Data Fig. 5B. **(D)** Schematic of the in vitro reconstitution of nascent VDAC3 post-translational insertion into the OM of purified mitochondria. Untagged VDAC3 is translated in RRL in the presence of ^35^S-labeled methionine and then incubated with mitochondria purified from either wildtype (wt) or FAM136A knockout (KO) human K562 cells. FAM136A KO was confirmed via western blot as shown in Extended Data Fig. 6B. Mitochondria were separated from any non-integrated translation products by centrifugation. To differentiate between the population of nascent VDAC3 in the IMS versus that in the OM, mitochondria were swollen using hypotonic buffer to perforate the OM without solubilizing it and treated with proteinase K (PK). Controls were used to demonstrate that under these conditions, any nascent substrates within the IMS become protease accessible, while the inner membrane (IM) remains intact and those integrated into the OM are protected (Extended Data Fig. 6A). The resulting samples representing the total mitochondrial population (IMS+OM+IM) and the protease protected fragment (OM+IM). We assume that VDAC3 is only inserted into the OM since the IM has no insertion machinery for β-barrel proteins. Therefore, we can neglect the contribution on the IM. **(E)** Autoradiography of a representative time course of ^35^S-methionine labelled VDAC3 insertion into purified mitochondria from K562 WT or FAM136A knockout (KO) cells. Samples are then swollen and subjected to PK treatment. Shown is both the mitochondrially integrated population (total) before PK treatment and the protease protected population after PK treatment (PK). **(F)** Quantification of the time course shown in (E). Shown is insertion of VDAC3 into K562 wt (white) or FAM136A KO (blue) cells over time. Data has been normalized to the maximum observed insertion (fraction inserted) and is displayed as the mean of three independent biological replicates ± SD, along with the non-linear fit of the data (see Materials and methods).

To determine whether this solubilizing effect was due to a direct physical interaction between FAM136A and VDAC3, we introduced a site-specific photo crosslinker in several positions in VDAC3. After exposure to UV light, we observed specific UV-dependent crosslinking between VDAC3 and FAM136A, consistent with chemical crosslinking experiments (**Fig. 3C, Extended Data Fig. 5B-C**). Because site-specific crosslinking can only occur for interactions <10Å ^41,42^, we concluded that FAM136A directly interacts with VDAC3.

We next tested if this direct interaction required other factors for substrate loading of VDAC3 onto FAM136A. To do this, we leveraged the PURE system, which contains only the purified recombinant *E. coli* translation machinery, but no chaperones or eukaryotic factors. Even in this minimal system, recombinant FAM136A was able to bind and immunoprecipitate nascent VDAC3 (**Extended Data Fig. 5D**). Therefore, no additional factors were required for FAM136A to capture nascent VDAC3. In summary, FAM136A is alone sufficient to act as a chaperone for unfolded, keeping them soluble in an aqueous environment similar to that in the IMS.

#### FAM136A mediates OM insertion of all three nascent VDAC paralogs from the IMS

Finally, and perhaps most importantly, if FAM136A functions as a chaperone, FAM136A should be required for VDAC integration into the OM. More specifically, if FAM136A maintains nascent VDACs in a soluble and insertion-competent state in the IMS, we expect it to be necessary to achieve the folded VDACs in the OM but not affect the total population that is imported into the mitochondria. To specifically query which step in VDAC biogenesis is affected by FAM136A depletion, we adopted an in vitro insertion assay to study the incorporation of VDAC3 into purified mitochondria (**Fig. 3D, Extended Data Fig. 6A**). Importantly, because we can use the untagged VDAC3 in these experiments, we can exclude any artifacts from tagging or the split-GFP system that was used in some earlier experiments. We translated and inserted VDAC3 into purified mitochondria from wildtype or FAM136A knockout cells and permeabilized the OM using hypotonic swelling. This method releases the IMS components while the inner membrane remains intact^43^. Further, it does not solubilize the outer membrane, ensuring the transmembrane regions of OM proteins remain protected from enzymatic digestion. We found that while the total amount of VDAC3 imported into mitochondria was not affected by loss of FAM136A, the amount that is protected after swelling and protease digestion is significantly decreased (**Fig. 3E, F**). We therefore concluded that in the absence of FAM136A, while VDAC3 is able to translocate into mitochondria equally efficiently across the OM, it is less competently folded and inserted into the OM. Validating that this observation is specifically due to loss of FAM136A, we demonstrated that expression of exogenous FAM136A rescues this effect for all three VDACs (**Extended Data Fig. 6B-C**).

In summary, FAM136A has all the features of an IMS-resident chaperone: it specifically and directly binds unfolded, nascent β-barrels in the IMS, keeping them soluble, and in a folding-competent state, thereby facilitating their integration into the OM.

### FAM136A confers substrate specificity via a hydrophobic groove lined with negative charges

Given that FAM136A is conserved in metazoa and plants, but not fungi, we next sought to determine what features of the mammalian VDACs might require an additional chaperone beyond the universally conserved small TIMs. Therefore, using VDAC3 as a model, we mapped the biophysical features of the interaction between FAM136A and its substrates.

We hypothesized that the interaction would be analogous to other membrane protein chaperones and rely on interaction between the VDACs’ exposed hydrophobic β-strands and FAM136A^44^. Indeed, we found that the interaction between VDAC3 and FAM136A in vitro was detergent sensitive (**Extended Data Fig. 7A**), which suggested that FAM136A must contain a hydrophobic surface for substrate binding. To identify this putative binding site, we examined the hydrophobicity of an AlphaFold model of the FAM136A dimer and identified a series of hydrophobic residues that together formed a continuous groove (**Fig. 4A; Extended Data Fig. 7B**)^45^. Mutations that disrupt this hydrophobic groove, but do not affect FAM136A expression or stability, rendered the chaperone unable to facilitate VDAC3 biogenesis in cells (**Fig. 4A; Extended Data Fig. 7B-C**). Consistent with these experiments, we also found that mutations that decreased the hydrophobicity of VDAC3’s β-strands disrupted FAM136A binding in vitro, while increasing its hydrophobicity had no effect (**Fig. 4B**). Based on these mutational analyses we reasoned that FAM136A interacted with the hydrophobic β-strands of its substrates via its prominent hydrophobic groove.

**Figure 4:**
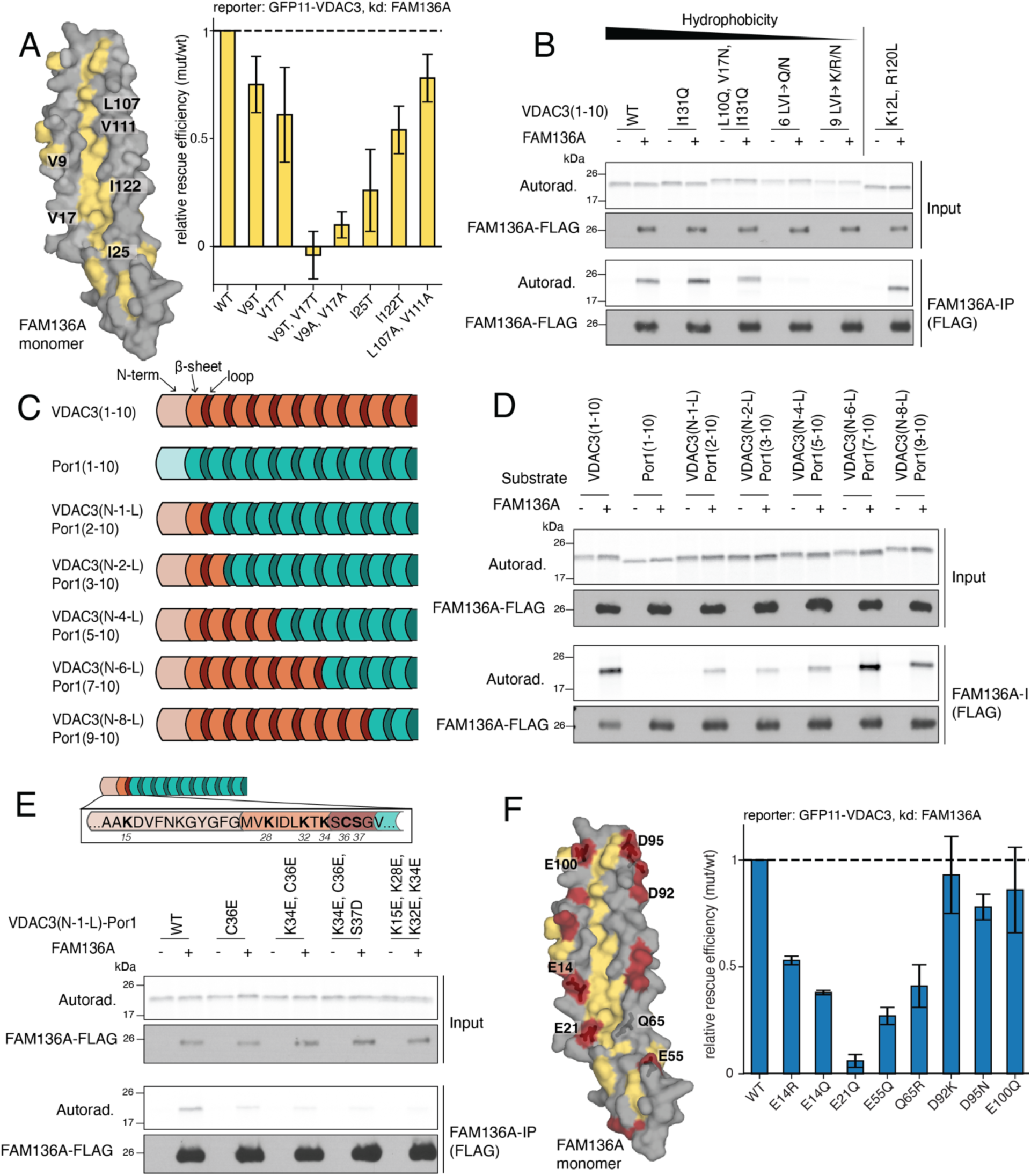
FAM136A interacts with VDAC3 via a hydrophobic groove lined by negative charges. **(A)** Mutational analysis to assess the role of the hydrophobic groove in FAM136A on VDAC3 biogenesis performed using the ratiometric fluorescent reporter system described in Fig. 1A. Wildtype (wt) FAM136A or the indicated mutants were transduced into FAM136A depleted K562 CRISPRi cells expressing GFP11-VDAC3. (Left) A space-filling representation of the predicted AlphaFold3^45^ model of a FAM136A monomer (alphafoldid: AF-Q96C01-F1-v6) is displayed in which the hydrophobic residues are colored in yellow and the residues that were mutated are indicated. (Right) Relative rescue efficiency of each FAM136A mutant on VDAC3 integration normalized to wt FAM136A rescue. Wt FAM136A rescue is set to 1 and the relative rescue efficiency of each mutant compared to wt is calculated and displayed as the mean ± SD of 2-3 independent biological replicates. Representative flow cytometry histograms are shown in Extended Data Fig. 7B and western blots assessing the expression of rescue constructs are shown in Extended Data Fig. 7C. **(B)** Analysis of the effects of altering the hydrophobicity of nascent VDAC3 on its ability to bind FAM136A in vitro. For simplicity, we used a truncated form containing only the first ten β - strands, VDAC3(1-10), which was mutated to successively decrease or increase its hydrophobicity. ^35^S-methionine labeled VDAC3(1-10) mutants were translated in RRL in the absence or presence of recombinant FAM136A-3xFLAG purified from *E. coli*. FAM136A was immunoprecipitated using anti-FLAG resin and eluted with 3xFLAG peptide. Co-purification of each mutant was analyzed by SDS-PAGE and autoradiography. Samples were also subjected to western blotting to ensure an equal amount of FAM136A-3xFLAG in both the input and elution. 6 LVI◊Q/N denotes: L10Q, V17N, I102Q, I131Q, V143N, L150Q. 9 LVI◊K/R/N denotes: L10K, V17N, V27N, L69K, L81K, I102R, I131R, V143N, L150K. **(C)** Schematic of the VDAC3-Por1(1-10) chimeric constructs used in (D). To further dissect the properties of the FAM136A-VDAC3 interaction, we leveraged the observation that Por1, the fungal homolog of the mammalian VDACs does not efficiently bind FAM136A in vitro (Extended Data Fig. 7D). Therefore, chimeras of VDAC3 and Porin1 could be used to determine the minimal sequence and its electrostatic properties required for FAM136A binding. For simplicity, we focused on a truncated form of VDAC3 and Por1 using only the first ten β-strands of each substrate. VDAC3 is shown in orange and Por1 in green, with their β-strands and intervening soluble loops displayed in light and dark shades, respectively. The chimeric constructs possess an increasing amount of VDAC3 β-sheets and loops on the Por1 background, progressively making Por1 more VDAC3-like. **(D)** Analysis of FAM136A binding of the nascent VDAC3-Por1 chimeras shown in (C) in vitro. The ^35^S-meyhionine labeled substrates were translated in RRL in the absence or presence of recombinant FAM136A-3xFLAG purified from *E. coli*. FAM136A was immunoprecipitated using anti-FLAG resin under native conditions and eluted with 3xFLAG peptide. Co-purification of each substrate was analyzed by SDS-PAGE and autoradiography. Samples were also subjected to western blotting to ensure an equal amount of FAM136A-3xFLAG in the input and elution. **(E)** To specifically probe the features that dictate FAM136A substrate binding, we leveraged the VDAC3(N-1-L)-Por1(2-10) chimera, which contains only a single FAM136A binding site composed of the N-terminus, first β-strand, and first loop of VDAC3. To test the role of charge in substrate binding, we generated a series of mutants to the VDAC3 sequence within the VDAC3(N-1-L)-Por1(2-10) chimera, in which positively charged lysine residues (K) were replaced with negatively charged glutamic acid (E). These ^35^S-methionine labeled constructs were translated in RRL in the absence or presence of recombinant FAM136A-3xFLAG purified from *E. coli*. FAM136A was immunoprecipitated under native conditions using anti-FLAG resin and eluted with 3xFLAG peptide. Co-purification of each mutant was analyzed by SDS-PAGE and autoradiography. Samples were also subjected to western blotting to ensure an equal amount of FAM136A-3xFLAG in the input and elution. **(F)** As in (A) to test the function of the conserved negatively charged residues that line the hydrophobic groove in FAM136A on VDAC3 biogenesis. (Left) A space-filling representation of the predicted AlphaFold3 model^45^ of a FAM136A monomer (alphafoldid: AF-Q96C01-F1-v6) is displayed in which the hydrophobic residues are colored in yellow and the negatively charged residues in red. (Right) Relative rescue efficiency of each FAM136A mutant on VDAC3 integration normalized to wt FAM136A rescue. Wt FAM136A rescue is set to 1 and the relative rescue efficiency of each mutant compared to wt is calculated and displayed as the mean ± SD of 2-3 independent biological replicates. Representative flow cytometry histograms are shown in Extended Data Fig. 9C and western blots assessing the expression of rescue constructs are shown in Extended Data Fig. 9D.

However, a hydrophobic interaction alone cannot explain FAM136A’s apparent substrate specificity or why the VDACs would require a distinct chaperone in the IMS. For example, if hydrophobicity alone dictated substrate binding to FAM136A, we would expect Porin1, the yeast VDAC homolog to also bind FAM136A. Despite the overall higher hydrophobicity of its β-strands (**Extended Data Fig. 8A-B**), we found that both full length Porin1 or a truncation of its first 10 β-strands failed to bind FAM136A in vitro (**Fig. 4D; Extended Data Fig. 7D**), suggesting that there must be other features that contribute to substrate specificity.

To dissect what these might be, we leveraged the observation that despite their nearly identical architecture, VDAC3 and Porin1 display distinct abilities to bind FAM136A. We therefore, systematically generated a series of VDAC3-Porin1(1-10) chimeras and tested them for binding to FAM136A in vitro (**Fig. 4C and Extended Data Fig. 7E**). We found that inclusion of the VDAC3 sequence spanning from its N-terminus through the first β-strand and loop was sufficient to confer FAM136A binding of our Porin1-chimera (**Fig. 4D, Extended Data Fig. 7F**). One potential interpretation of this result is that the N-terminus of VDAC3 represents a specific FAM136A binding site. However, inclusion of increasing amounts of VDAC3 (both β-strands and intervening loops) in the Porin1 chimera resulted in a commensurate increase in binding to FAM136A. We therefore instead concluded that features of VDAC3 confer FAM136A binding along multiple sites along the nascent protein, but not within a single unique binding site.

To further dissect the requirements for substrate binding, we used our identified minimal FAM136A binding site—the N-terminus of VDAC3 through the first loop—fused to Porin1. We note that within this region of VDAC3 there are several positively charged residues that are not presented in Porin1, which overall is enriched for negative charge relative to all three VDAC paralogs (**Extended Data Fig. 8A and C**). To test if charge may contribute to FAM136A binding, we generated a series of mutants within this minimal binding chimera. Overall, we found that introduction of negative charge prevented binding to FAM136A in vitro, but mutations to neutral residues did not affect binding (**Fig. 4E; Extended Data Fig. 9A**).

Electrostatic analysis of the FAM136A dimer showed that the identified hydrophobic groove is lined by a prominent series of negative charges, which is conserved across metazoans and plants (**Fig. 4F; Extended Data Fig. 9B; Fig. 6B**). Indeed, mutations that disrupted the charge of these residues but do not affect FAM136A expression or stability, resulted in loss of function in cells (**Fig. 4F; Extended Data Fig. 9C-D**). Taken together, we concluded that FAM136A binds its substrates via a hydrophobic groove lined with negatively charged residues. Enrichment of positively charged amino acids in the VDACs appeared to encourage recruitment to FAM136A, while enrichment of negatively charged residues, such as in Porin1, prevent binding presumably through electrostatic repulsion.

### FAM136A’s chaperone function plays a unique role in the IMS for both β-barrels and a subset of IM α-helical proteins

Given the specific features of VDAC3 that confer FAM136A dependence, we next sought to understand how FAM136A fit in the larger context of the IMS chaperone network. First, we performed a genetic modifier screen to identify factors that have enhanced or diminished phenotypes upon depletion of FAM136A^46^. Since FAM136A functions in the IMS, we focused our analysis on other IMS localized factors. We found that the disaggregase CLPB and its chaperone, HAX1^47–49^—both of which have been previously shown to be localized to the IMS and potentially implicated in FAM136A function^34,35^—become slightly more significant hits in our FAM136A knockdown vs wildtype cells (**Fig. 5A; Extended Data Fig. 10A**). Depletion of both FAM136A and CLPB or HAX1 had an additive effect on VDAC3 biogenesis, suggesting they function in parallel or partially redundant pathways (**Extended Data Fig. 10B-C**). We therefore proposed that CLPB’s disaggregase function may solubilize VDAC3 in the IMS, facilitating its integration into the OM via a mechanism distinct and potentially unrelated to FAM136A activity.

**Fig. 5:**
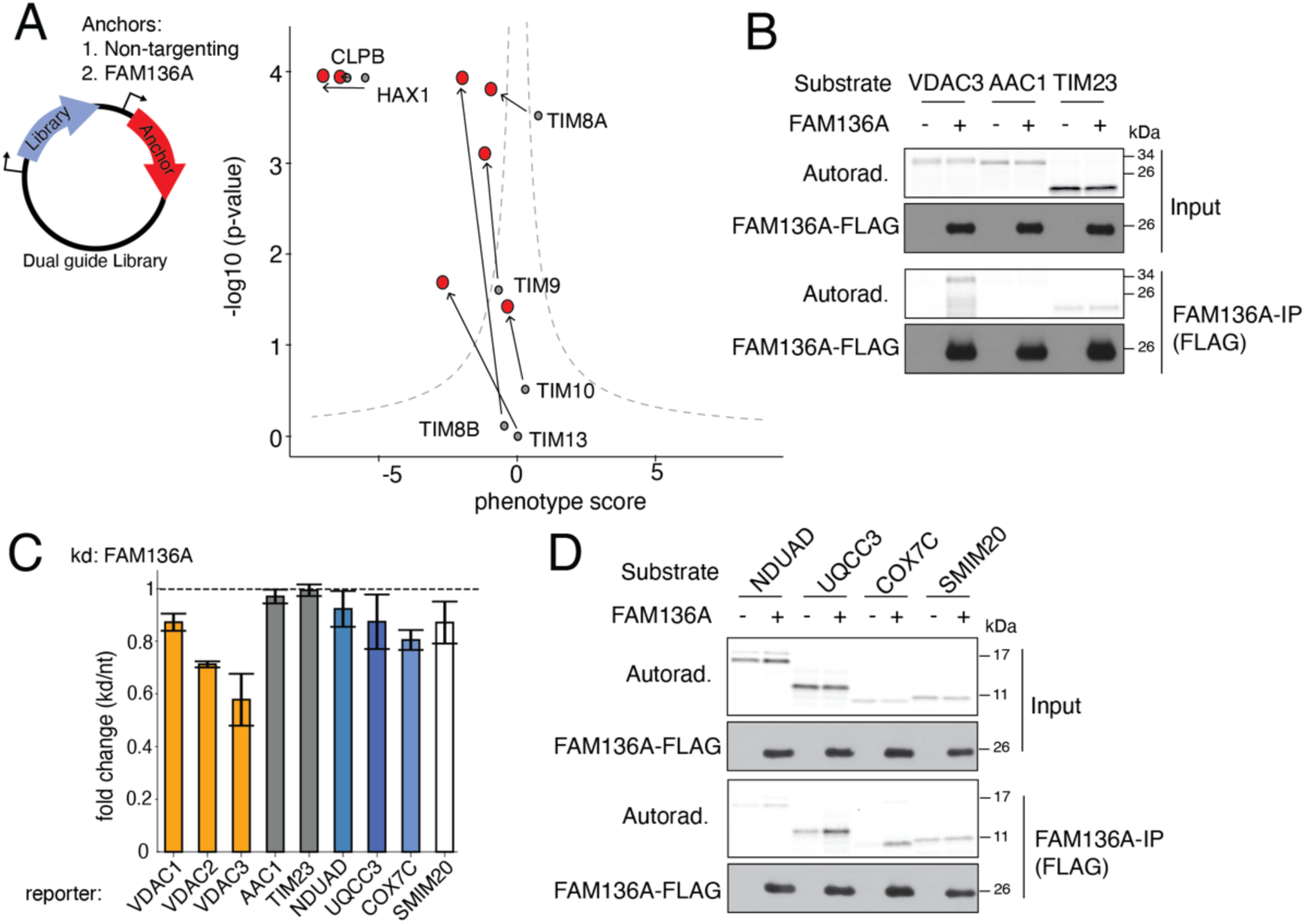
FAM136A plays a non-overlapping function to the TIM chaperones in biogenesis of both OM β-barrels and IM a-helical proteins. **(A)** To understand how FAM136A fits in the larger context of the IMS chaperone network genetic modifier screen to identify factors that have enhanced or diminished phenotypes on VDAC3 biogenesis in human cells upon depletion of FAM136A was performed ^46^. For this, a genome wide CRISPRi dual-screen using the VDAC3 reporter depicted in (**A**) was used. (Left) Schematic of the dual-guide library design. Expression of the CRISPRi TOP5 v2 sgRNA library driven by a mU6 promoter while a fixed anchor guide (either non-targeting control or FAM136A) is driven by a hU6 promoter. (Right) CRISPRi cells stably expressing GFP11-VDAC3 along with the normalization marker (RFP) were transduced with a modified TOP5 v2 sgRNA library harboring either a non-targeting control or FAM136A as genetic anchor. Following eight days of depletion, cells with altered GFP:RFP ratios were isolated using FACS and deep sequenced to identify enriched sgRNAs in the top (increased GFP:RFP) and bottom (decreased GFP:RFP) populations. Depicted is a differential volcano plot showing the phenotype score versus the -log of the Mann-Whitney p values. Shown are the disaggregase CLPB and its chaperone HAX1 as well as the small TIM proteins in the nt control-anchored screen in grey and in the FAM136A-anchored screen in red. Arrows indicate the change between the two screens by starting in the position in the nt-anchored screen and ending in the position in the FAM136A-anchored screen. Volcano plots with all genes displayed are shown in Extended Data Fig. 10A. **(B)** Binding of the canonical TIM substrates AAC1 and TIM23 to FAM136A in vitro. The ^35^S-meyhionine labeled AAC1 or TIM23 were translated in rabbit reticulocyte lysate (RRL) in the absence or presence of recombinant FAM136A-3xFLAG purified from *E. coli*. FAM136A was immunoprecipitated using anti-FLAG resin and eluted with 3xFLAG peptide. Co-purification of AAC1 or TIM23 was analyzed by SDS-PAGE and autoradiography. Samples were also subjected to western blotting to ensure an equal amount of FAM136A-3xFLAG in the input and elution. **(C)** FAM136A kd affects VDACs but not canonical TIM substrates AAC1 and TIM23 in cells. GFP11-tagged VDAC1, VDAC2, VDAC3, AAC1, TIM23 or inner mitochondria membrane located single TM reporters signal change was assessed in K562 CRISPRi cells that constitutively expresses GFP1-10 in the IMS as well as a nontargeting control (nt) or FAM136A knockdown sgRNA as described in Figure 1A. Shown is the fold change of the GFP:RFP ratio of the reporters in kd over nt control cells (kd/nt) as mean ± SD, n=3. **(D)** Binding of the inner mitochondria membrane localized single TM proteins to FAM136A in vitro. The ^35^S-meyhionine labeled IM proteins were translated in the PURExpress system (NEB) in the absence or presence of recombinant FAM136A-3xFLAG purified from *E. coli*. FAM136A was immunoprecipitated using anti-FLAG resin and eluted with 3xFLAG peptide. Co-purification of IM proteins was analyzed by SDS-PAGE and autoradiography. Samples were also subjected to western blotting to ensure an equal amount of FAM136A-3xFLAG in the input and elution.

Given FAM136A’s analogous role to the small TIM chaperones, we next examined whether perturbing proteostasis in the IMS, through depletion of FAM136A might alter TIM activity on VDACs biogenesis. Comparison of our genetic modifier and traditional CRISPRi screen suggested that while depletion of TIM8B or TIM13, which function as a heterohexameric chaperone^50,51^, had no phenotype on VDAC3 alone (**Fig. 1B**), in the absence of FAM136A their depletion becomes more significant (**Fig. 5A; Extended Data Fig. 10A**). This suggested that TIM8B/TIM13 may be partially able to chaperone nascent VDAC3, but only when its primary chaperone is unavailable.

Conversely, FAM136A does not efficiently bind the canonical TIM substrates AAC1 or TIM23^19^ under in vitro conditions where it successfully captures VDAC1-3 (**Fig. 5B**). Concordant with our in vitro assays, we find that FAM136A knockdown does not affect TIM23 and AAC1 biogenesis in cells (**Fig. 5C**). Therefore, FAM136A plays a unique and non-overlapping role to the small TIMs for β-barrel biogenesis in the IMS.

However, β-barrel proteins are not the only nascent substrates that require chaperoning within the IMS. The small TIMs also play a role in the biogenesis and targeting of nuclear-encoded α-helical subunits to the inner membrane. Indeed, depletion of FAM136A affected the biogenesis of several single-spanning representative members of each of the ETC complexes (**Fig. 5C**)^34,35^. We concluded that these effects were specific because many IM substrates remained unaffected by FAM136A depletion (**Extended Data Fig. 10D**), excluding a general dysregulation of mitochondrial protein biogenesis or the ETC. Finally, these α-helical substrates directly bound to FAM136A in vitro (**Fig. 5D**), and their biogenesis was not affected by the small TIM proteins in cells (**Extended Data Fig. 10E**). Together, these data support a specific role for FAM136A in chaperoning the VDACs to the OM and a subset of small single-pass α-helical inner membrane proteins.

## DISCUSSION

In summary, our data describe a critical step in biogenesis of mitochondrial membrane proteins in human cells. For β-barrels, after their synthesis in the cytosol, the nascent, extended VDACs must be recognized and targeted to the mitochondria (**Fig. 6A**). Because their mitochondrial targeting signal is localized within the C-terminus, this must occur post-translationally after release from the ribosome^52–54^. Translocation across the OM occurs via the TOM complex, which requires the nascent protein to remain in an extended, unfolded state to fit through the translocation pore of TOM40^55^.

**Fig. 6:**
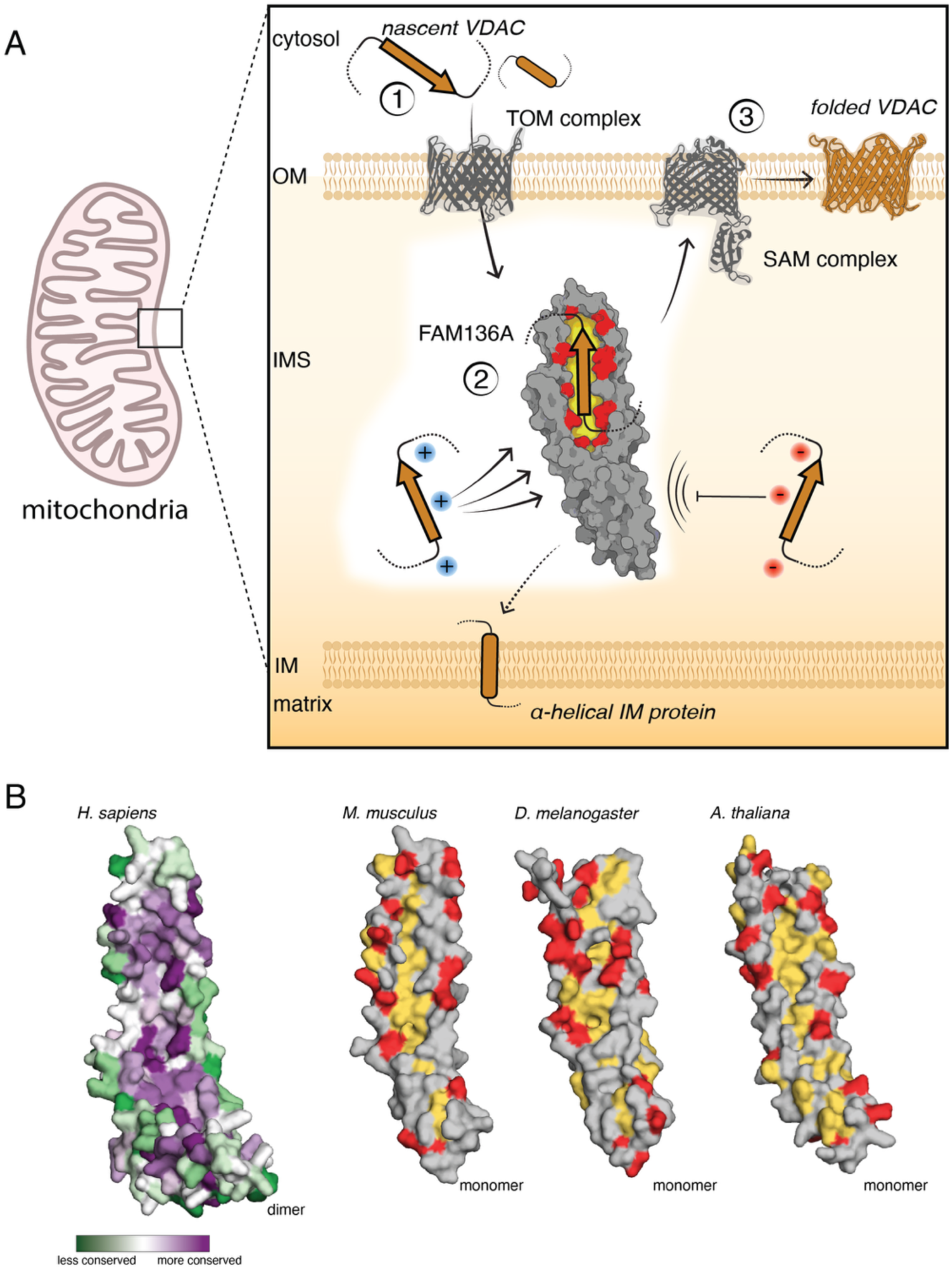
A model of the conserved FAM136A chaperone function. **(A)** Model of FAM136A chaperone function. (1) Nascent VDAC or hydrophobic α-helical inner membrane protein is targeted to the OM and translocates via the TOM complex into the IMS. (2) Here, the nascent, unfolded VDAC protein is chaperoned by FAM136A via binding to its hydrophobic groove. The negative charges lining the groove provide substrate specificity selecting for positively charged substrates such as VDAC3 and against negative charges on substrates as in the case of Porin1. (3) VDAC is then inserted into the OM via the SAM complex where it assumes its final fold while hydrophobic α-helical inner membrane proteins are targeted and inserted to the IM. **(B)** FAM136A functional features are conserved. Left: Conservation of each residue of human FAM136A was calculated using the ConSurf software^66–68^. A space-filling representation of the predicted AlphaFold3 model of a FAM136A dimer (alphafoldid: AF-0000000065888360-v1) is displayed in which the value of conservation of each residue is indicated by colors with purple indicating highly conserved residues and green indicating non-conserved residues. Particularly residues in the and around the hydrophobic groove are highly conserved. Right: Space-filling representation of the predicted AlphaFold3 models of FAM136A monomers from three different species (mouse – *M. musculus* (uniprotid: Q9CR98); fruit fly – *D. melanogaster* (uniprotid: Q9W3A7); plant – *A. thaliana* (uniprotid: Q5Q0I6)) are displayed in which the hydrophobic residues are colored in yellow, and the negatively charged residues are colored in red.

Upon exposure to the IMS, our data suggests that the nascent β-strands of VDAC1, 2, and 3 are then captured by the symmetric FAM136A dimer. While it remains to be seen if there is a coordinated hand-over from the TOM complex or even interaction of FAM136A with the IMS-exposed extension of TOM40, similar to what has been shown for the small TIMs^56^, our data suggest that FAM136A can directly bind nascent β-strands as they are emerging into the IMS. FAM136A contains a hydrophobic groove of ∼1000 Å^2^, which is sufficient to accommodate 1-2 β-strands and our studies show that five β-strands is sufficient for VDAC binding to FAM136A. Conserved negative charges surrounding the hydrophobic groove contribute to substrate selectivity, primarily by excluding substrates with intervening acidic sequences, or stabilizing binding of substrates enriched for basic residues. All three VDAC paralogs are de-enriched for negative charges, potentially explaining their binding and recruitment by FAM136A (**Extended Data Fig. 8A**). The role of charge has been previously shown to play a crucial role in several aspects of membrane protein biogenesis across all kingdoms of life. For example, electrostatic repulsion has previously been described to regulate insertion of IM proteins via the TIM23 complex. Here negatively charged substrates are repelled by negatively charged residues in TIM17, arresting them during their import and allowing for lateral release into the IM ^57^. Similarly, electrostatic interactions also contribute to the binding between the bacterial chaperone Skp and unfolded outer membrane proteins (uOMPs) in the periplasm, the ancestral compartment to the IMS^58,59^.

The combination of the hydrophobic groove and negative charged residues make FAM136A binding sufficient to maintain the solubility of the unfolded VDACs, preventing non-specific interactions and off-pathway aggregation while the hydrophobic β-sheets transit through the aqueous environment of the IMS. Our data suggest that under certain conditions, multiple FAM136A dimers can bind the intact 19-strand VDAC1,2, and 3 in the IMS. However, the relative kinetics of translocation into the IMS versus integration into the OM will dictate the number of β-strands exposed to the IMS that must be captured and shielded by FAM136A.

Genetic interaction analysis suggests that FAM136A carries out this function in a parallel pathway to the disaggregase CLPB (which itself relies on a chaperone for assembly, HAX1^48^), which can also bind unfolded VDAC and may also facilitate its solubilization within the IMS. While CLPB and HAX1 levels had previously been reported to be modestly decreased by FAM136A depletion^34,35^, our data suggests the effects on VDAC biogenesis after FAM136A depletion are not based on the loss of these disaggregase components. Conversely, our data supports direct binding of FAM136A to nascent VDACs via hydrophobic and electrostatic interactions. Further FAM136A sequentially decorates the translocated β-sheets to keep nascent VDACs in an import-competent state and facilitates hand-off to the SAM complex for templated integration into the OM.

FAM136A also appears to play an analogous role in chaperoning a subset of IM proteins within the IMS. These include small (<150 amino acids), single-spanning subunits of the ETC or the mitochondrial translation regulation assembly intermediate of cytochrome c oxidase (complex IV) (MITRAC) in the case of SMIM20, which are characterized by a hydrophobic α-helical TM. Many of the IM subunits identified as putative FAM136A substrates are also enriched for positive charges, which may suggest a coherent mechanism for substrate selection across both β-barrel and α-helical membrane proteins. A role for FAM136A in the biogenesis of several IM subunits could in part explain earlier experiments connecting FAM136A to ETC complex expression^34,35^.

This function as an IMS chaperone, contributing to the biogenesis of all three VDAC paralogs as part of this physiologically important, highly expressed, class of β-barrel proteins and some IM proteins, and particularly required for nascent VDAC3, provides a biochemical explanation for its essentiality in cells and previously reported pleotropic phenotypes when depleted^39,60^.

The role of FAM136A may be critically important in cells with high metabolic needs, including inner ear hair cells. Due to the unique signaling properties of these cells, they have exceptionally high basal ATP requirements and they cannot regenerate after damage, and therefore may be particularly sensitive to changes in VDAC3 levels, the paralog that protects cells from accumulation of ROS and regulates respiration^61^. The nonsense mutation within FAM136A associated with hearing loss in Meníere’s disease fails to localize to the IMS^35,60^. Loss of FAM136A chaperone function and the resulting defects in VDAC and ETC biogenesis in these unique cells provides a biochemical explanation for the disease phenotype.

However, we must further consider the role of FAM136A within the context of the network of biogenesis and quality control factors that populate the IMS. The IMS is a highly specialized cellular compartment ^62^. It has an unusual ionic and redox environment, and though overall protein concentrations have not been rigorously measured, because of its very small volume estimates of individual proteins such as cytochrome C suggest local concentrations as high as 1 mM^63^. Transiting through this space is also a mixture of nascent proteins headed to several locations: soluble proteins localized for the matrix, α-helical membrane proteins targeted to the IM, and finally β-barrel proteins that must be integrated into the OM.

Therefore, chaperones within the IMS have two distinct functions. First, they must keep nascent proteins soluble, shielding hydrophobic stretches within this crowded environment and second, they must selectively target proteins to distinct locations. Here we show that FAM136A is capable of the former and systematically define its biophysical features that allow it to specifically capture its substrates. We have made initial strides towards a deeper understanding of the latter. Indeed, FAM136A appears to be required beyond the universally conserved TIM proteins primarily in organisms with expanded mitochondrial proteomes. This may reflect both the expanded chaperone repertoire required to capture increased sequence and biophysical diversity, but also specialization, essential to minimize competition between many nascent substrates to ensure targeting specificity.

Collectively, our results suggest that the complexity of the IMS must mimic that of the bacterial periplasm. In bacteria, a network of chaperones and proteases together regulate uOMPs biogenesis: chaperoning nascent β-barrels to the membrane, facilitating proper folding into the OM by direct handoff to insertion machinery, and finally triaging misfolded and aggregated substrates to degradation to prevent toxicity ^6^. We envision the IMS chaperone network must contain similar features, that we are now in a position to define at the molecular level. We propose that, similar to periplasmic chaperones, FAM136A and the TIMs must therefore function in concert to create a network of protein biogenesis and quality control factors capable of providing directionality and accurate targeting to ensure mitochondrial protein homeostasis and function.

## ACKNOWLEDGEMENTS

We thank T. Stevens for thoughtful discussion and input on the manuscript. We thank: the Caltech Flow cytometry facility; the Proteome Exploration Laboratory at Caltech for mass spectrometry; the Caltech Millard and Muriel Jacobs Genetics and Genomics Laboratory; and the Caltech Biological Imaging Facility. **Funding.** This work was supported by an NIGMS K99 (5K99GM151478) to AG. RMV is a Howard Hughes Medical Institute Freeman Hrabowski Scholar. **Competing interests.** RMV is an equity holder and consultant for Gate Biosciences.

## METHODS

### Plasmids

#### Protein sequences

Endogenous sequences used in this study for in vitro and cell-based assays were sourced from UniprotKB/Swiss-Prot and included: voltage dependent anion channel 1 (VDAC1; P21796), voltage dependent anion channel 2, (VDAC2; P45880), voltage dependent anion channel 3 (VDAC3; Q9Y277), translocase of inner mitochondrial membrane 8A (TIM8A; O60220), translocase of inner mitochondrial membrane 8B (TIM8B; Q9Y5J9), translocase of inner mitochondrial membrane 9 (TIM9; Q9Y5J7), translocase of inner mitochondrial membrane 10 (TIM10; P62072), translocase of inner mitochondrial membrane 13 (TIM13; Q9Y5L4), translocator protein (TSPO; P30536), single stranded DNA binding protein 1 (SSBP1; Q04837), family with sequence similarity 136, member A (FAM136A; Q96C01), translocase of outer mitochondrial membrane 40 (TOMM40; O96008), translocase of inner mitochondrial membrane 23 (TIM23; O14925), solute carrier family 25 member 4 (AAC1; P12235), synaptojanin-2 binding protein (OMP25/SYNJBP; P57105-1), suppressor of potassium transport defect 3 (CLPB/SKD3, Q9H078), HCLS1-associated protein X-1 (HAX1, O00165).

#### General plasmid backbones

The 2nd generation lentiviral packaging plasmid pCMV-VSV-G was a gift from Bob Weinberg (Addgene #8454). The 2nd generation lentiviral packaging plasmid psPAX2 was a gift from Didier Trono (Addgene #12260). The pHAGE2 lentiviral transfer plasmid was a gift from Magnus A. Hoffmann and Pamela Bjorkman. The dual guide lentiviral vector pJR103 was a gift from Jonathan Weissman (Addgene #187242). The SFFV-tet3G backbone was used for K562 cell expression during CRISPRi screens^69^.

#### Reporter constructs in CRISPRi cells

For the in-cell experiments assessing reporter stability changes as described in **Fig. 1A**, the protein coding sequences were cloned into the dual color split reporter system (RFP-P2A-GFP11) which has been previously described^30,70,71^. The reporter constructs were cloned into a mammalian expression lentiviral backbone containing a UCOE-EF-1α promoter or a 3’ WPRE element^72^ (Addgene #135448). Note that the mCherry variant of RFP was used in all instances, but the simpler nomenclature of RFP is used in the text and figures. For complementation with the GFP1-10 system, the GFP11 tag (RDHMVLHEYVNAAGIT) was appended to the appropriate terminus of the indicated protein as determined by predicted topology or based on previous data^30^. For targeting GFP1-10 to the intermembrane space of the mitochondria, the targeting sequence from LACTB (residues 1–68)^73^ was appended to the N-terminus of GFP1-10 as described previously^30^. For some reporter constructs that reside in the inner mitochondrial membrane or the matrix (**Fig. 1G**), the GFP1-10 was targeted to the mitochondrial matrix via appending the targeting sequence from COX4 to the N-terminus of GFP1-10 as previously described^64^. To check mis-localization of FAM136A to the ER, the GFP1-10 was targeted to the ER lumen via a the human calreticulin signal sequence preceding a GFP1-10-KDEL^74–76^.

#### VDAC3 reporter construct in K562 CRISPRi cells for CRISPRi screens

The VDAC3 reporter used for the CRISPRi screen shown in **Fig. 1C** was cloned into the dual color reporter system described above with GFP11 and a 10 amino acid linker on the N-terminus of VDAC3 and integrated into an indicuble doxycycline-inducible tet3G backbone^69^.

#### CRISPRi knockdown guides

Programmed single and dual guides were generated to assay depletion of one or two genes using the CRISPRi system. Single sgRNA were generated by annealing oligonucleotides (Integrated DNA Technologies, Coralville, IA), which were introduced into a lentiviral plasmid with a mouse U6 promoter for the sgRNA and an EF-1α promotor preceding a Puro-P2A-BFP sequence digested with BstXI/BlpI (Addgene #84832). BFP was excised in certain sgRNA variants where BFP was instead needed to indicate expression of a second cDNA, such as FAM136a rescue experiments (**Fig. 4A,F; Extended Data Fig. 4A; Extended Data Fig. 6B; Extended Data Fig. 7B; Extended Data Fig. 9C**). Dual sgRNA were generated by annealing two pairs of oligonucleotides (Integrated DNA Technologies, Coralville, IA) and cloning into a lentiviral mouseU6-sgRNA1-humanU6-sgRNA2 backbone^46,77^. The sgRNA guide sequences used are shown in **Table S2.**

The shRNA against TOM40 was purchased from Sigma Aldrich Mission shRNA with the targeting sequence CAAAGGGTTGAGTAACCATTT.

#### Constructs for in vitro translation

Constructs for expression in rabbit reticulate lysate (RRL) were cloned into the SP64 vector (Promega, Madison, WI). For transcription, the DNA template was PCR amplified using the P2long and P3 or Gblock_rev primers (**Table S3**) and the Q5^®^ High-Fidelity 2X Master Mix (#M0492, NEB, USA)^78^. Constructs for expression in the PURE system^79,80^ were based on the T7 PURExpress plasmid provided by New England Biolabs.

#### Constructs for recombinant protein expression from E. coli

Constructs for human FAM136A and CLPB expression in *E. coli* were cloned into the pQE plasmid (Qiagen, Valencia, CA) downstream of a His14-SUMO^Eu1^ tag. FAM136A was fused with a C-terminal 3xFLAG tag and CLPB was fused with a C-terminal ALFA tag to be used in immunoprecipitations (IP) assays.

#### Constructs for recombinant protein expression from Expi293F cells

Constructs for generating human stable cell lines for recombinant protein expression used the pHAGE2 plasmid (gift from Magnus A. Hoffmann and Pamela Bjorkman) for lentiviral integration into the Expi293F cell line. FAM136A was C-terminally fused with an ALFA tag downstream of a CMV promoter.

All individual plasmids are listed in **Table S1** and are available on request.

### Antibodies

The following antibodies were used in this study:

#### Primary antibodies

SAMM50 (ab133709, Abcam, UK), tubulin (T9026, Sigma-Aldrich, USA), TOM40 (sc-365467, Santa Cruz Biotech, USA), TIM8A (11179-1-AP, Proteintech, USA), TIM8B (144-61221-20, RayBiotech, USA), TIM9 (sc-101285, Santa Cruz Biotech, USA), TIM10 (11124-2-AP, Proteintech, USA), TIM13 (11973-1-AP, Proteintech, USA), FAM136A (CSB-PA836190LA01HU, Cusabio, USA); HAX1 (ab137613, Abcam, UK), CLPB (15743-1-AP, Proteintech, USA), VDAC1/3 (ab14734, Abcam, UK), VDAC2 (ab37985 Abcam, UK).

#### Secondary antibodies used for immunoblotting

goat anti-mouse-HRP (172-1011, Bio-Rad, USA) and anti-rabbit-HRP (170-6515, Bio-Rad, USA).

#### Others

The ALFA tag was detected by coupling HRP to an ALFA nanobody as described previously^30^; FLAG-HRP (A8592, Sigma-Aldrich, USA, RRID:AB_439702; 1:10,000)

### Cell culture and cell line generation

#### General cell culture conditions

K562 cells were grown in RPMI-1640 with 25 mM HEPES, 2.0 g/L NaHCO3, and 0.3 g/L L-glutamine supplemented with 10% FBS fetal bovine serum (FBS; S11150, Bio-Techne, USA) or Tet System Approved FBS for the CRISPRi screen, 100 units/mL penicillin, and 100 μg/mL streptomycin. Cells were maintained at a confluency between 0.25 × 10^6^ –1 × 10^6^ cells/mL. HEK293T cells were grown in Dulbecco’s Modified Eagle Medium (DMEM; 11965092, Thermo Scientific, USA) supplemented with 10% FBS and 1X L-glutamine.

RPE1 cells were grown in Dulbecco’s Modified Eagle Medium/Nutrient Mixture F-12 (DMEM/F-12; 11320033, Thermo Scientific, USA) supplemented with 10% FBS and 1X L-glutamine. Expi293F cells were grown in Expi293^TM^ expression medium and maintained at a confluency between 0.5 × 10^6^ –2 × 10^6^ cells/mL. K562, HEK293T and RPE1 cells were grown at 37°C, 5% CO_2_, and Expi293F cells were grown at 37 °C, 8% CO_2_ while shaking at 125 rpm.

#### VDAC3 CRISPRi screen cell line

To generate the VDAC3 screen cell line used in the CRISPRi screen shown in **Fig. 1C**, K562-dCas9-BFP-KRAB Tet-On cells^69^ stably expressing LACTB-GFP1-10 cells were infected with lentivirus delivering the tet3G-mCherry-P2A-GFP11-10aa-VDAC3 construct and sorted into 96-well plates as single cell clones using a Sony Cell Sorter (SH800S). Cells were grown in RPMI-1640 with 25 mM HEPES, 2.0 g/L NaHCO3, and 0.3 g/L L-glutamine supplemented with 20% Tet System Approved FBS, 100 units/mL penicillin, and 100 μg/mL streptomycin. After expansion, correct cell lines were confirmed by induction with doxycycline (100 ng/mL) for 48 h.

#### FAM136A knockout cell line

To generate the FAM136A monoclonal knockout cell line, K562 CRISPRi cells with LACTB(GFP1-10) were nucleofected with a FAM136A targeting guide in the pX458 backbone (#48138, Addgene) using the Lonza SF Cell Line 96-well Nucleofector Kit (V4SC-2096). The pX458 backbone was adapted to express a sgRNA targeting FAM136A [GGCTTTGGTCACCAGTGAGC] as well as GFP. Two days following nucleofection, GFP-positive cells indicating incorporation of the guide were sorted into 96-well plates. After several weeks of expansion, loss of FAM136A was confirmed for each monoclonal cell line by immunoblotting.

#### FAM136A-ALFA Expi293F stable expression cell line generation

Expi293F cells were transduced by adding 2.5 mL of FAM136A-ALFA-P2A-BFP-3xFLAG lentivirus to 15 million cells along with 8 µg/mL final concentration of polybrene in a volume of 30 mL in a 125-mL cell culture flask. After 8h, media was exchanged to remove lentiviral particles and prevent cell clumping. BFP positive cells were isolated as a polyclonal population using a SONY SH800S cell sorter (Sony Biotechnology, USA).

### Endogenous tagging

Tagging of VDAC3 at its respective endogenous loci in K562 CRISPRi cells was performed as previously described^81^. Briefly, to endogenously tag VDAC3 with mCherry-P2A-GFP11 positioned at the N-terminus, a protospacer within exon three was cloned into pX458 (aacgtactgtgacctaggaa). A vector encoding mCherry-P2A-GFP11 and 800 bp of homology on either side of the cut site of the VDAC3 locus was ordered from TWIST Biosciences. The resulting vector was transduced, and RFP positive cells were sorted into 96-well plates as single cell clones using a Sony Cell Sorter (SH800S). Cells were grown in RPMI-1640 with 25 mM HEPES, 2.0 g/L NaHCO3, and 0.3 g/L L-glutamine supplemented with 20% FBS, 100 units/mL penicillin, and 100 μg/mL streptomycin. After expansion, correct cell lines were confirmed by FACS analysis of RFP and GFP positive cells indicating incorporation of the endogenous tag.

### Lentivirus production

Lentivirus was generated by co-transfecting HEK293T cells seeded the previous day at 0.2 × 10^6^ cells/mL and 2.5 mL in a 6-well plate with two packaging plasmids (pCMV-VSV-G and ΔpPAX, Addgene #8454) and the desired transfer plasmid using TransIT-293 transfection reagent (Mirus). 48 h after transfection, the supernatant was collected, aliquoted and frozen. In all instances, virus was rapidly thawed prior to transfection. Virus for the genome wide CRISPRi screen was also generated using this method.

### Ratiometric fluorescent reporter assays to test effects of individual depletion of genes

Throughout the manuscript we rely on depletion of individual genes to assess the effect of loss of specific factors on a fluorescent reporter. These reporters serve two purposes. First, because the reporters express a substrate (e.g. VDAC3-GFP11) and a normalization marker (e.g. RFP) from a single open reading frame separated by a viral 2A ribosomal skipping site, changes in GFP:RFP ratio necessarily reflect changes that occur post-translationally. Second, because the reporter only expresses the GFP11 fragment while the GFP1-10 fragment is stably expressed in the IMS (or some cases the matrix), the observed GFP signal reflects successful integration of the newly synthesized protein into mitochondria.

#### Single and dual knock-down reporter assay in K562 CRISPRi cells and K562 CRISPRi VDAC3-GFP11 cells

For reporter assays probing mitochondrial reporter proteins, as shown in **Fig. 1B,E,G; Fig. 5C; Extended Data Fig. 1A-B**; **Extended Data Fig. 2B-D; Extended Data Fig. 10B,E-F**), K562 cells stably expressing ZIM3 KRAB-dCas9-P2A-BFP from a UCOE-SFFV promoter^69^ as well as either LACTB-GFP1-10 (IMS targeted) or COX4-GFP11 (matrix targeted) were used. The sgRNA guides were on a PURO-P2A-BFP backbone and the reporter sequences were on a dual color reporter system (RFP-P2A-GFP11). Lentivirus containing sgRNAs targeting a gene of interest or a non-targeting control were prepared and spinfected into K562 CRISPRi cells at density 0.25 × 10^6^ cells/mL along with 8 µg/mL final concentration of polybrene. Cells containing the knockdown guide were selected via a single dose puromycin treatment (3 μg/mL) 48 h after spinfection for 72 h. On day five, reporter lentivirus was spinfected at a density of 0.25 × 10^6^ cells/mL along with 8 µg/mL final concentration of polybrene. Cells were analyzed on day eight on an Attune NxT Flow Cytometer (Thermo Scientific, USA) and cells were gated for BFP (integration of the sgRNA guide) and RFP (integration of the reporter virus). Since the K562 cells with endogenously tagged RFP_P2A_GFP11_VDAC3 cells as shown in **Fig. 1F** are already expressing the fluorescent reporter endogenously, these cells did not get spinfected with reporter lentivirus on day 5 but were cultured until analysis on day 8.

#### Reporter assay in K562 FAM136A KO cells

For fluorescent reporter assays in FAM136A KO cells as shown in **Extended Data Fig. 6B**, reporter lentivirus was spinfected into FAM136A KO cells at a density of 0.25 × 10^6^ cells/mL and 8 µg/mL final concentration of polybrene. Cells were analyzed 72 h after spinfection on an Attune NxT Flow Cytometer (Thermo Scientific, USA) and cells were gated for RFP (integration of the reporter virus).

#### Rescue assay in K562 CRISPRi cells

To probe the effect of re-introducing FAM136A or FAM136A mutants after knockdown as shown in **Fig. 4A, F; Extended Data Fig. 4A; Extended Data Fig. 7B; Extended Data Fig. 9C**, K562 cells stably expressing ZIM3 KRAB-dCas9-P2A-BFP from a UCOE-SFFV promoter^69^ as well as LACTB-GFP1-10 (IMS targeted) were used^30^. Here, colorless (no BFP) sgRNA guides expressed from a backbone along with a puromycin resistance marker a were used. The reporter sequences were expressed using the ratiometric fluorescent reporter system (RFP-P2A-GFP11) and the rescue constructs were expressed as either untagged or ALFA-tag fusions from a P2A_BFP backbone. Lentivirus containing sgRNAs targeting a gene of interest or a non-targeting control were prepared and spinfected into K562 CRISPRi cells at density 0.25 × 10^6^ cells/mL using 8 µg/mL final concentration of polybrene. Cells containing the knockdown guide were selected via a single dose of puromycin treatment (3 μg/mL) 48 h after spinfection for 72 h. On day five, reporter and rescue construct lentivirus were spinfected as described above. Cells were analyzed on day eight on an Attune NxT Flow Cytometer (Thermo Scientific, USA) and cells were gated for BFP (integration of the rescue construct virus) and RFP (integration of the reporter virus).

#### Flow cytometry to analyze fluorescence reporter assays

K562 cells were analyzed directly from cultures using an Attune NxT Flow Cytometer (Thermo Scientific, USA). Generated flow cytometry data was then analyzed by Python (version 2.7) using the FlowCytometryTools package^82^. For knockdown reporter assays, cells were gated first for live cells; then for BFP positivity, reflecting expression of the sgRNA guide or cDNA rescue construct; and finally for RFP positivity, reflecting integration and expression of the reporter construct. The GFP:RFP ratio was calculated via the median of the GFP and RFP signals. The relative change (KD/NT or KO/WT) of each reporter was calculated by taking the ratio of the GFP:RFP intensities between that of the respective knockdown (KD) or knockout (KO) cells and the non-targeting (NT) or wild type (WT) control cells (Equation 1). Relative change is displayed in bar plots showing the mean ± standard deviation.

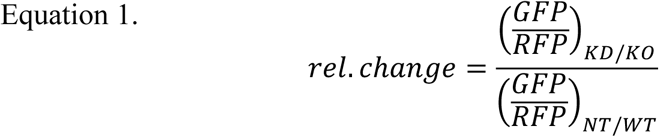

The rescue efficiency of FAM136A mutants after FAM136A KD was calculated by normalizing the GFP/RFP to a FAM136A WT rescue construct control. (Equation 2). Relative rescue efficiency is displayed in bar plots showing the mean ± standard deviation.

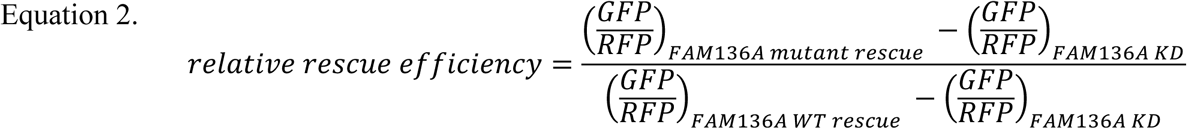

Histograms, charts, and plots were generated using Python (version 2.7).

### Western blotting of knockdown cells

After eight days of knockdown, ∼500,000 – 1,000,000 cells were harvested and washed three times with PBS. Cells were lysed in lysis buffer (50 mM HEPES pH 7.5, 200 mM NaCl, 1 mM DTT, 1x cOmplete^TM^ EDTA-free Protease Inhibitor Cocktail (Roche)) with 1% glyco-diosgenin (GDN, Anatrace) and centrifuged to remove debris. The total protein concentration in the supernatant was measured via A_280_, and 20 μg protein/lane were loaded onto an SDS-PAGE gel. Proteins were then transferred onto a nitrocellulose membrane and blocked with 5 % milk powder in Tris-buffered saline with Tween^®^ 20 (TBST). Membranes were incubated with primary antibody in 5% milk in TBST over night at 4 °C. After washing three times 5 minutes with TBST, membranes were incubated with secondary antibody for one hour at room temperature.

### CRISPRi screen

The genome-scale CRISPR interference (CRISPRi) screen, as shown in **Fig. 1C**, was performed as described previously^30,32,33,46,64^. The human dual-sgRNA library based on the hCRISPRi-v2 compact library (5 sgRNAs per gene, Addgene pooled library #83969) was designed in previous studies ^46^. This allows for the simultaneous delivery and selection of a fixed sgRNA and a second randomized guide, comprised of a genome-wide library (hCRISPRi-v2 compact library), with a single transduction^46^. Here we used a non-targeting control as the fixed sgRNA in the VDAC3 screen shown in **Fig. 1C**. We also used a FAM136A targeting sgRNA for a second screen as shown in **Fig. 5A; Extended Data Fig. 10A**. The library was spinfected into 400 million K562-CRISPRi-Tet-ON-((LACTB)-GFP1-10)-(tet-RFPP2A-GFP11-VDAC3) cells at multiplicity of infection (MOI) < 1 (percentage of transduced cells 48 h after infection as measured by BFP positive cells was around 30%). To ensure that the culture was maintained at an average coverage of more than 1000 cells per sgRNA, cells were diluted daily to 0.5 × 10^6^ cells/mL and maintained in 1 L of media in 1 L spinner flasks (Bellco, SKU: 1965-61010). Cells containing an sgRNA knockdown guide were selected 48 h after spinfection with 1 μg/mL puromycin for three days. Cells were allowed to recover from the puro treatment for 36 h, induced with 100 ng/mL doxycycline for 36 h and finally sorted on a FACS AriaII Fusion Cell Sorter on day 8.

During sorting, cells were gated for BFP positivity (indicating a guide-positive cell), as well as RFP positivity (successfully induced). Cells were sorted based on the GFP:RFP ratio of this final gated population. Roughly 30 million cells with either the highest (30%) or lowest (30%) RFP:GFP ratio were collected, washed with PBS, pelleted and flash-frozen. Genomic DNA was extracted using the Nucleospin Blood XL kit (Takara Bio, #740950.10) and amplified by index PCR with barcoded primers. The resulting guide library (∼349 bp) was purified using SPRIbeads (SPRIselect Beckman Coulter #B23318) and sequencing was performed using an Illumina HiSeq2500 high throughput sequencer. Sequencing reads were aligned to the CRISPRi v2 library sequences, counted and quantified as described previously^32,33^. Generation of negative control genes and calculation of phenotype scores and Mann–Whitney p-values was performed as described previously ^32,33^. Gene-level phenotypes and counts are available in source file 1.

### Cell fixation and confocal microscopy of mitochondrial morphology

Circular cover glass (72222-01, Electron Microscopy Sciences, USA) was sterilized, placed in the bottom of each 12-well plate, and coated by incubating with 1 mL of 0.01 mg/mL poly-D-lysine hydrobromide (P6407, Sigma-Aldrich, USA) for 1 h at room temperature. Coated cover glass was washed 3 times with sterilized water and dried overnight at 37 °C. To study mitochondrial morphology by confocal microscopy (**Fig. 1H**), RPE1 cells were transduced with lentivirus containing the indicated sgRNAs at a final concentration of 8 µg/mL polybrene. After 48 h, cells were treated with 8 µg/mL puromycin for two days and then recovered in regular media for another 3 days. Approximately 60,000 cells were plated per well in a 12-well plate with a coverslip bottom prepared as described above. To visualize mitochondria, cells were incubated with DMEM/F-12 containing 150 nM MitoTracker DeepRed FM (Thermo Fisher, M22426) at 37 °C for 45 min. Cells were then rinsed once in fresh media and twice in ice-cold 1x PBS. Cells were fixed in 4% paraformaldehyde (PFA; 15714, Electron Microscopy Sciences) in 1x PBS for 15 min at room temperature. Post fixation, cells were washed three times with ice-cold 1x PBS and permeabilized in 0.1% Triton X-100 in PBS for 5 min at room temperature. Samples were finally stained with Samples were stained with 1 µg/mL DAPI (MBD0015, Sigma-Aldrich, USA) in 1x PBS for 5 min and washed 2 times with 1x PBS, followed by 2 washes with water. To mount samples onto microscope slides, each stained cover glass was inverted onto 7 µL of SlowFade^TM^ Gold Antifade Mountant (S36936, Invitrogen, USA) added on a glass microscope slide. Cover slips were sealed and secured in place on microscope slides with nail polish. Prepared microscope slides were stored at room temperature overnight to dry and stored at 4 °C until imaging. Imaging was performed on a Zeiss LSM 980 with Airyscan2 Laser Scanning Confocal microscope with 63x oil-immersion objective.

### Mass spectrometry sample preparation of FAM136A from mitochondria immunoprecipitations

To determine interaction partners of FAM136A in mitochondria as shown in **Fig. 2A**, crude mitochondrial isolation and anti-ALFA immunoprecipitation of exogenously expressed FAM136A-ALFA from Expi293F cells were conducted as described previously^83,84^. The control sample for background binding of FAM136A immunoprecipitation in **Fig. 2A** was wt Expi293F cells not overexpressing FAM136A-ALFA. Briefly, cells were centrifuged for 5 min at 300x g, and the pellets were washed once in homogenization buffer (210 mM mannitol, 70 mM sucrose, 5 mM HEPES-KOH pH 7.5, 1 mM EDTA, cOmplete^TM^ EDTA-free Protease Inhibitor Cocktail (Roche)). The weight of the resulting cell pellet was calculated, and 2 mL of homogenization buffer was added per 0.5 g of pellet. After resuspension, cells were incubated on ice for 10 min, before a 2 mL glass Dounce homogenizer with a tight-fitting pestle was used to lyse cells using ∼30 passes. The lysate was centrifuged at 1300x g for 5 min and the supernatant was transferred to a clean tube to remove nuclei and unlysed cells. To pellet mitochondria, the supernatant was then centrifuged at 11,000x g for 10 min.

The mitochondrial pellets were incubated in solubilization buffer (50 mM HEPES-NaOH pH 7.5, 200 mM NaCl, 2 mM MgOAc_2_, 1% [w/v] glycol-diosgenin [GDN, GDN101, Anatrace, USA], cOmplete^TM^ EDTA-free Protease Inhibitor Cocktail (Roche), 1 mM DTT) for 30 min at 4 °C while rotating. Solubilized mitochondria were clarified by centrifugation at 18,000 rpm in an SS-34 rotor in a Sorvall RC6+ Superspeed centrifuge for 30 min at 4 °C. At the same time, biotinylated His14-Avi-SUMOStar-tagged anti-ALFA nanobody (See below for preparation method in “Recombinant protein expression and purification from *E. coli*<u>”</u>) was incubated with Pierce^TM^ Streptavidin Magnetic Beads (88817, Thermo Scientific, USA) head-over-tail in wash buffer (50 mM HEPES-NaOH pH 7.5, 200 mM NaCl, 2 mM MgOAc_2_, 0.0053% [w/v] glycol-diosgenin [GDN, GDN101, Anatrace, USA], 1 mM DTT) for 30 min at 4 °C. After this immobilization, beads were washed and incubated with 50 mM HEPES pH 7.5 containing 100 µM biotin on ice for 5 min to occupy unbound biotin binding sites. Then, beads were washed in solubilization buffer twice before being incubated with clarified lysate head-over-tail for 1.5 hr at 4 °C. After incubation, beads were washed 3 times in wash buffer. Proteins were eluted by addition of 0.3 µM SUMOStar protease and incubated for 30-45 min at 4 °C while shaking. Eluted protein samples were precipitated by adding 1:10 volume of 100% trichloroacetic acid (TCA), followed by a 10 min incubation on ice before being pelleted at maximum speed in a benchtop centrifuge at 4 °C. Protein pellets were washed with ice-cold acetone twice and then air dried at room temperature. TCA-precipitated protein pellets were dissolved in 8 M urea prepared in 50 mM HEPES pH 8.0 and samples were reduced by incubating with 4 mM Tris(2-carboxyethyl)phosphine hydrochloride (TCEP) (20490, Thermo Scientific, USA) for 20 min at 37 °C while shaking at 750 rpm. Samples were alkylated by incubating with 12 mM 2-chloro-acetamide (CAA) (ICN15495580, MP Biomedicals) for 15 min at 37 °C and then digested with 2 ng/µL Lysyl Endoproteinase (Lys-C) (125-05061, Wako Chemicals) for 4 h at 37 °C with shaking. Samples were diluted with 50 mM HEPES pH 7.5 to a final concentration of 2 M urea, before being digested overnight with 0.6 ng/µL Trypsin (90057, Thermo Scientific, USA). Samples were desalted using the Pierce^TM^ C18 Spin Columns (89870, Thermo Scientific, USA) as described by the manufacturer’s instructions. After being eluted off the desalting columns, samples were lyophilized before mass spectrometry analysis.

### LC-MS/MS analysis for FAM136A interaction partners

Data was acquired in data dependent acquisition (DDA) mode on an Orbitrap Eclipse Tribrid mass spectrometer (Thermo Fisher Scientific, USA) coupled to a Vanquish Neo UHPLC system (Thermo Fisher Scientific, USA). Peptides were separated on an Aurora UHPLC Column (25 cm × 75 μm, 1.7 μm C18, AUR3-25075C18-TS, Ion Opticks) with a flow rate of 0.35 μL/min and ionized at 1.8 kV in the positive ion mode. The active gradient was composed of 6% B (3.5 min), 6-25% B (41.5 min), 25-40% B (15 min), 40-98% B (2 min) and 98% B (5min); mobile phase A is 2% acetonitrile (ACN, Fisher Scientific, A9554) and 0.2% formic acid (FA, Fisher Scientific, A11750) in water (Optima LC-MS grade, Fisher Scientific, W6212), and mobile phase B is 80% ACN and 0.2% FA in water. MS1 spectra were collected in the Orbitrap at the resolution of 120,000 from 375 to 1,600 m/z with automatic gain control (AGC) target of 250% and a maximum injection time of 50 ms. MS2 scans were acquired in the ion trap using fast scan rate on precursors with 2-7 charge states and quadrupole isolation mode (isolation window: 1.2 m/z) with higher-energy collisional dissociation (HCD, 30%) activation type. Dynamic exclusion was set to 15 s and low/high mass tolerance was 5ppm. Ion transfer tube temperature was 300°C and the S-lens RF level was set to 30.

MS RAW files were searched against the Uniprot reviewed human database (swiss-prot, UP000005640) using the MaxQuant (2.3.0.1). Trypsin/P was set as the digestion enzyme, allowing a maximum of 2 missed cleavages. Oxidation / +15.995 Da (M) and acetylation / +42.011 Da (N-term) were set as dynamic modifications, and carbamidomethylation / +57.021 Da (C) was fixed modification. Label-free quantification (LFQ) and Intensity-based absolute quantification (iBAQ) were performed to quantify protein abundance across samples. The mass spectrometry proteomics data have been deposited to the ProteomeXchange Consortium via the PRIDE^85^ partner repository with the dataset identifier PXD080641. They can be accessed via log in to the PRIDE website using the following details: **Project accession:** PXD080641, **Token:** E1WgIPn4o2RE. Data used for generating Fig. 2A can be found in source data file 2.

### Anti-ALFA immunoprecipitation of FAM136A, TIM9 and TIM10 from Expi293F cells

To compare interaction partners of FAM136A, TIM9 and TIM10 in cells (**Fig. 2B**), each of the three proteins was overexpressed and immunoprecipitated from Expi293F cells with a C-terminally fused ALFA tag. Anti-ALFA immunoprecipitation of exogenously expressed protein was performed as described previously^84^. Briefly, cells were harvested by centrifugation and washed with ice-cold 1x PBS. Cell pellets were resuspended in solubilization buffer (50 mM HEPES-NaOH pH 7.5, 200 mM NaCl, 2 mM MgOAc_2_, 1% [w/v] glycol-diosgenin [GDN, GDN101, Anatrace, USA], cOmplete^TM^ EDTA-free Protease Inhibitor Cocktail (Roche), 1 mM DTT) for 30 min at 4 °C while rotating. Solubilized lysates were clarified by centrifugation at 18,000 rpm in an SS-34 rotor in a Sorvall RC6+ Superspeed centrifuge for 30 min at 4 °C. Simultaneously, biotinylated His14-Avi-SUMOStar-tagged anti-ALFA nanobody was incubated with Pierce^TM^ Streptavidin Magnetic Beads (88817, Thermo Scientific, USA) head-over-tail in wash buffer (50 mM HEPES-NaOH pH 7.5, 200 mM NaCl, 2 mM MgOAc_2_, 0.0053% [w/v] glycol-diosgenin [GDN, GDN101, Anatrace, USA], 1 mM DTT) for 30 min at 4 °C. After this immobilization, beads were washed and incubated with 50 mM HEPES pH 7.5 containing 100 µM biotin on ice for 5 min to occupy unbound biotin binding sites. Then, beads were washed in solubilization buffer twice before being incubated with clarified lysate head-over-tail for 1.5 h at 4 °C. After incubation, beads were washed 3 times in wash buffer. Proteins were eluted by addition of 0.3 µM SUMOStar protease and incubated for 30-45 min at 4 °C while shaking. Eluted samples were analyzed using SDS-PAGE.

### Mitochondrial fractionation and immunoprecipitation of FAM136A from the IMS

To determine interaction partners of FAM136A specifically in the IMS (**Fig. 2C**), we isolated an exogenously expressed FAM136A under native conditions from the IMS. To do this, we used the FAM136A-ALFA Expi293F stable expression cell line for crude mitochondria preparation. Isolation of mitochondria was conducted as described previously^83^. Briefly, 200 mL of Expi293F cells overexpressing FAM136A-ALFA were centrifuged for 5 min at 300x g, and the pellet was washed once in homogenization buffer (210 mM mannitol, 70 mM sucrose, 5 mM HEPES-KOH pH 7.5, 1 mM EDTA, cOmplete^TM^ EDTA-free Protease Inhibitor Cocktail (Roche)). The weight of the resulting cell pellet was calculated, and 2 mL of homogenization buffer was added per 0.5 g of pellet. After resuspension, cells were incubated on ice for 10 min, before a 2 mL glass Dounce homogenizer with a tight-fitting pestle was used to lyse cells using ∼30 passes. The lysate was centrifuged at 1300x g for 5 min, and the supernatant was transferred to a clean tube to remove nuclei and unlysed cells. This step was repeated twice, before mitochondria were pelleted by centrifugating at 11,000x g for 10 min. Pelleted mitochondria were washed twice in isolation buffer (210 mM mannitol, 70 mM sucrose, 5 mM HEPES-KOH pH 7.5, 1 mM EDTA) and the final pellet was resuspended in 15-20 µL of isolation buffer. Prior to downstream use, protein concentration was measured using a Bradford assay.

To isolate only the population of FAM136A from the IMS, purified mitochondria were subjected to swelling ^43^. Briefly, mitochondria were pelleted for 5 min at 11,000 × g at 4 °C, resuspended in hypotonic buffer (10 mM HEPES pH 7.2, 1 mM EDTA) to a final protein concentration of 1 mg/mL, and incubated for 25 min at 4 °C. The resuspension was then centrifuged for 10 min at 18,000 × g at 4 °C, and the supernatant was pelleted again at 45,000 rpm in an Optima^TM^ MAX-XP table top ultracentrifuge (Beckman-Coulter) for 15 min at 4 °C. After centrifugation, the supernatant was mixed with same volume of 2x wash buffer (100 mM HEPES-NaOH pH 7.5, 400 mM NaCl, 4 mM MgOAc_2_, 2 mM DTT). Western blotting for a representative protein from the OM (MTCH2) was performed to determine successful isolation of IMS (**Fig. 2C**). Meanwhile, biotinylated His14-Avi-SUMOStar-tagged anti-ALFA nanobody was incubated with Pierce^TM^ Streptavidin Magnetic Beads (88817, Thermo Scientific, USA) head-over-tail in wash buffer (50 mM HEPES-NaOH pH 7.5, 200 mM NaCl, 2 mM MgOAc_2_, 1 mM DTT) for 20 min at 4 °C. After this immobilization, beads were washed and incubated with 50 mM HEPES pH 7.5 containing 100 µM biotin on ice for 5 min to occupy unbound biotin binding sites. Then, beads were washed in solubilization buffer twice before being incubated with clarified lysate head-over-tail for 1.5 h at 4 °C. After incubation, beads were washed 3 times in wash buffer. Proteins were eluted by addition of 0.3 µM SUMOStar protease and incubated for 30-45 min at 4 °C while shaking. Eluted sample was analyzed by SDS-PAGE and western blotting.

### Recombinant protein expression and purification from *E. coli*

For in vitro immunoprecipitations in **Fig. 2D**; **Fig. 3B-C; Fig. 4B,D,E; Fig. 5B,D; Extended Data Fig. 4D; Extended Data Fig. 5B-D; Extended Data Fig. 7A,D,F; Extended Data Fig. 9A** the following recombinant proteins were expressed and purified from *E. coli* with slightly different strategies.

#### Expression and purification of recombinant FAM136A-3xFLAG

To express recombinant FAM136A-3xFLAG protein, plasmids were transformed into BL21 (DE3) *E. coli* expression strain. A single colony was picked and inoculated into 20 mL of Luria-Bertani (LB) medium supplemented with 50 µg/mL kanamycin and incubated at 37 °C while shaking at 220 rpm overnight. The next day, 5 mL of overnight culture was inoculated into 1 L of Super Broth (SB) medium supplemented with 50 µg/mL kanamycin incubated at 37 °C while shaking at 200 rpm. When OD_600_ reached 0.6-0.8, cells were induced with 0.5 mM isopropyl-β-D-thiogalactopyranoside (IPTG) and grown overnight at 18 °C while shaking. Cells were harvested by centrifugation, resuspended in 10 mL of lysis buffer (50 mM HEPES-NaOH pH 7.5, 200 mM NaCl, 10% [v/v] glycerol), flash frozen in liquid nitrogen, and stored at −80 °C for later use.

After thawing, the cells were resuspended in 30 mL with lysis buffer and supplemented with 20 mM imidazole, 5 mM β-mercaptoethanol and cOmplete^TM^ EDTA-free Protease Inhibitor Cocktail (Roche). Cells were lysed by sonication, and the soluble fraction was isolated by centrifugation at 18,000 rpm and 4 °C for 30 min using an SS-34 rotor (Beckman-Coulter). Cleared lysate was loaded onto a gravity flow column (Bio-Rad) containing 5 mL of Ni-NTA agarose (Qiagen). Ni-NTA resin was then washed with 80 mL of wash buffer (50 mM HEPES-NaOH pH 7.5, 200 mM NaCl, 20 mM imidazole, 5 mM β-mercaptoethanol), 50 mL of high salt buffer (50 mM HEPES-NaOH pH 7.5, 1 M NaCl, 50 mM imidazole, 5 mM β-mercaptoethanol), and another 140 mL of wash buffer. Before eluting, Ni-NTA resin was finally washed with 20 mL of elution buffer (50 mM HEPES-NaOH pH 7.5, 150 mM NaCl, 10 mM imidazole, 10% [v/v] glycerol, 5 mM β-mercaptoethanol). Protein was eluted by incubating with 2 mL of elution buffer supplemented with 0.5 µM SENP^EuB^ protease (Addgene #149333, see below for preparation method) at 4 °C for 1 h, and then another 20 mL of elution buffer was applied to the gravity flow column to ensure thorough elution of cleaved protein. Eluted protein was concentrated to 500 µL using a 10K MWCO concentrator (Millipore-Sigma) and injected onto a Superdex 200 increase 10/300 GL size exclusion column (Cytiva) pre-equilibrated in SEC buffer (50 mM HEPES-NaOH pH 7.5, 200 mM NaCl, 2 mM MgOAc_2_, 1 mM DTT). Protein-containing fractions were collected, concentrated, flash frozen and stored at −80 °C for later use.

#### Generation of anti-ALFA nanobodies

The biotinylated His14-Avi-SUMOStar-tagged anti-ALFA nanobody was prepared as previously described^84^. Proteins were expressed in Rosetta-gami 2 *E. coli* strain and purified using Ni-NTA agarose beads, followed by in vitro biotinylation with purified BirA.

#### Generation of SENP^EuB^ protease

SENP^EuB^ protease used in the above purifications were prepared as previously described^84^. Proteins were expressed in NEBExpress *E. coli* strain and purified using Ni-NTA agarose beads.

### In vitro translation in rabbit reticulocyte lysate (RRL)

For immunoprecipitations in **Fig. 2D**; **Fig. 3B-C**; **Fig. 4B,D,E; Fig. 5B,D; Extended Data Fig. 4D; Extended Data Fig. 5B-D; Extended Data Fig. 7A,D,F; Extended Data Fig. 9A** in vitro translation reactions were performed as previously described with the following modifications^78^. In brief, the protein of interest is translated in rabbit reticulocyte lysate (RRL) supplemented with either 0.5 µM bait protein (recombinant FAM136A-3xFLAG or CLPB-ALFA) or buffer (50 mM Tris-HCl pH 7.5, 200 mM NaCl, 2 mM MgOAc_2_, 1 mM DTT) in the presence of ^35^S-methionine to allow for tracking by autoradiography following SDS-PAGE. DNA templates for in vitro transcription were made by PCR using primers within the SP6 promoter (5’ end) and after the stop codon (3’ end) as described above. Transcription reactions were carried out using an in-house transcription mix (T1)^78^ supplemented with 0.02X (v/v) RNAsin (N251, Promega, USA) and 0.02X (v/v) SP6 polymerase (M0207L, New England Biolabs, USA) and 5 ng/μL PCR product. Transcription reactions were incubated at 37°C for 2 h and then used either directly in a translation reaction or flash frozen. Translation reactions were carried out by combining transcripts with an in-house translation mix (CT2, ^78^), radioactive ^35^S-methionine (NEG709A005MC, Perkin Elmer, USA) and bait proteins to a final volume of 50 µL. Reactions were incubated at 32 °C for 30 min and treated with 1 mM puromycin at 32 °C for 10 min immediately following translation to fully release nascent proteins from the ribosome.

### In vitro translation in PURE

For immunoprecipitations in **Fig. 5D; Extended Data Fig. 5D**, in vitro translation reactions were carried out using the PURExpress In vitro Protein Synthesis Kit (NEB, E6800L). In brief, translations were performed in 10 µL reactions supplemented with 1 µM bait protein (recombinant FAM136A-3xFLAG) or buffer (50 mM Tris-HCl pH 7.5, 200 mM NaCl, 2 mM MgOAc_2_, 1 mM DTT) in the presence of ^35^S-methionine. Reactions were incubated at 32°C for 2 h and treated with 1 mM puromycin at 32 °C for 10 min immediately following translation to fully release nascent proteins from the ribosome.

### Immunoprecipitation after in vitro translation in RRL and PURE

For immunoprecipitations, 20 µL of FLAG beads (50% v/v in FLAG-IP buffer, Millipore-Sigma) were added directly to the translation samples and reactions were incubated head-over-tail for 1.5 h at 4 °C. After immobilization, samples were washed three times with FLAG-IP buffer (50 mM HEPES-NaOH, pH 7.5, 130 mM KOAc, 2 mM MgOAc_2_) and proteins were eluted with 0.2 mg/mL 3xFLAG peptide diluted in elution buffer (50 mM HEPES-NaOH, pH 7.5, 130 mM KOAc, 2 mM MgOAc_2_, 0.05% TritonX-100). Samples were boiled at 95 °C for 5 min and analyzed on SDS-PAGE followed by autoradiography.

For **Extended Data Fig. 7A**, to study the nature of the interaction between VDAC3 and FAM136A, TritonX-100 was used to test if the binding could still happen in the presence of detergent. In vitro transcription, translation and FLAG-IP were conducted as described above, except for that after immobilization, FLAG beads were washed three times with FLAG-IP buffer supplemented with increasing amount of TritonX-100 (0.1%, 0.5% and 1%), and proteins were eluted directly with 2.5X sample buffer. Samples were boiled at 95 °C for 5 min and analyzed on SDS-PAGE followed by autoradiography.

### Fractionation of in vitro translation via sucrose gradient

Sucrose gradients as shown in **Fig. 3B-C; Extended Data Fig. 5A-C** were prepared by layering 40 μL of 25%, 20%, 15%, 10% and 5% sucrose in phosphate buffered saline (PBS) buffer into ultracentrifuge tubes and incubating for one hour at 4 °C. Translation reactions (20 μL) were layered on top of sucrose gradient and centrifuged for 2 h at 55,000 rpm in an Optima^TM^ MAX-XP table top ultracentrifuge (Beckman-Coulter). Fractions of 20 μL were collected and analyzed on SDS-PAGE followed by autoradiography or used for crosslinking.

#### Chemical crosslinking of VDAC3 and FAM136A

To chemically crosslink VDAC3 to FAM136A as shown in **Extended Data Fig. 5C**, fractionated translation reactions were incubated with either 250 μM DSS in DMSO or DMSO for 30 min at room temperature. The reaction was then quenched with 20 mM Tris/HCl, pH 7.5 for 15 min on ice. Samples were denatured with 1% SDS and further analyzed via denaturing FLAG-IP as described above. Briefly, denatured samples were diluted in 1 mL FLAG-IP buffer (50 mM HEPES, pH7.5, 130 mM KAc, 2 mM MgAc, 5% TritonX-100). 20 μL pre-equilibrated FLAG beads (50% v/v in FLAG-IP buffer) were added and samples were incubated head-over-tail for 1.5 h at 4 °C. After immobilization, samples were washed three times with FLAG-IP buffer and FLAG-FAM136A-bound proteins were eluted with the addition of SDS sample buffer. Samples were boiled at 95 °C for 5 min and analyzed on SDS-PAGE followed by autoradiography.

### Site-specific crosslinking of VDAC3 and FAM136A

To crosslink VDAC3 to FAM136A as shown in **Fig. 3A-C; Extended Data Fig. 5B**, VDAC3 constructs harboring amber stop codon mutations at position (VDAC3*) were created and translated in the presence of Bpa-tRNA to incorporate the non-canonical amino acid Bpa into the protein. Bpa can then be photochemically activated via UV light and crosslinks to any other protein within 10 Å^41,42^.

#### Purification of T7 RNA-Polymerase (T7 RNAP)

The T7 RNAP sequence (p6XHis-T7(P266L)), was a gift from Anna Pyle (Addgene #174866). T7 RNAP was expressed in BL21(DE3) cells, induced with 1 mM IPTG and grown for 5 h at 37°C and 200 rpm shaking. The cells were harvested and lysed in in Buffer A (50 mM Tris pH=8.0, 100 mM NaCl, 10 mM imidazole, 5 mM beta mercaptoethanol, supplemented with Roche’s protease inhibitor tablet, 5% glycerol) via sonication and purified via nickel agarose column beads (QIAgen, Ni-NTA Agarose). Samples were washed with Buffer B1 (50 mM Tris pH=8.0, 1000 mM NaCl, 40 mM imidazole, 5 mM beta mercaptoethanol, 5% glycerol) and Buffer B2 (50 mM Tris pH=8.0, 100 mM NaCl, 40 mM imidazole, 5 mM beta mercaptoethanol, 5% glycerol) and eluted with Buffer C (50 mM Tris pH=8.0, 100 mM NaCl, 500 mM imidazole, 5 mM beta mercaptoethanol, 5% glycerol). Elution was then diluted 12-fold with Buffer EQ (20 mM Tris pH=8.0, 1 mM EDTA, 5 mM beta mercaptoethanol) and further purified via anion exchange column (Bio Rad, Macro-Prep High Q). Samples were washed with 100 EQ : 0 HS (20 mM Tris pH=8.0, 1 M NaCl, 1 mM EDTA, 5 mM beta mercaptoethanol, 10% glycerol), and 95 EQ : 5 HS with 5 CVs each. Protein was eluted with 35% HS content, concentrated (Amicon® Ultra 30K) and rebuffered with Buffer GF (50 mM Tris pH=8.0, 100 mM NaCl, 1 mM EDTA, 5 mM beta mercaptoethanol, degassed). Finally, the rebuffered protein further purified via size exclusion (Cytiva Superdex, S200 3.2/300), again concentrated (Amicon® Ultra 30K) and re-buffered with Buffer Y (50 mM Tris pH8, 100 mM NaCl, 1 mM EDTA, 0.1% Triton X, 10 mM beta mercaptoethanol, 50% glycerol).

#### In-vitro transcription of Bpa-tRNA(UAG)

For transcription, the DNA template (pLB144 a gift from the Hegde lab-pRMV457, **Table S1**) was PCR amplified using the LB130F and LB130R (**Table S3**) and the Q5^®^ High-Fidelity 2X Master Mix (#M0492, NEB, USA).

tRNA was transcribed with in-house purified T7 RNAP in 220 mM Tris-HCl, 11 mM spermidine, 137.5 mM MgCl2, 0.55 % Triton-X100 supplemented with 7.5 mM ATP, UTP, GTP and CTP, as well as 5 mM DTT, inorganic pyrophosphatase (NEB) and RNAsin for 16 h at 37 °C. After incubation, 0.5 uL of TURBO DNAse (Invitrogen) was added per 10 uL or reaction and incubated for 15 min at 37 °C. RNA was extracted by adding 3 volumes of TriZol and 0.6 volumes of chloroform, incubation at room temperature for 5 min and centrifugation at max speed for 10 min at room temperature. Then the aqueous phase was collected and 0.5 volumes of chilled isopropanol was added and incubated at room temperature for 10 min followed by centrifugation at 7000 xg, 4 °C for 5 min. The supernatant was aspirated and the RNA was dried for at least 15 min and resuspended in water to 2000 ng/uL.

#### Purification of Bpa aminoacyl synthetase

For expression, BAS (pLB152 a gift from the Hegde lab-pRMV456) was grown from a BL21 (DE3) glycerol stock in 150 mL LB media supplemented with 100 µg/mL carbenicillin over night at 37°C. The overnight culture was then diluted to an OD_600_ of 0.1 in 4 L SB media with 100 µg/mL carbenicillin and induced with 1 mM ITPG at OD_600_ of 0.4. The shaker temperature was reduced to 25°C and cells were grown over night at 200 rpm. The next day, cells were harvested via centrifugation at 4500 xg for 20 min and two pellets of 20 g were flash frozen and stored at −80°C. On the day of purification, one pellet was thawed on ice and lysed in in binding buffer (25 mM Hepes, 70 mM NH_4_Cl, 30 mM KCl, 7 mM MgCl_2_, 10% glycerol, 10 mM imidazole, 4 mM beta-mercaptoethanol, supplemented with Roche’s protease inhibitor tablet) via sonication. Lysate was cleared by pelleting membranes at 18000 rpm at 4°C for 40 min and the supernatant was incubated while rotating with equilibrated Ni-NTA resin for 1 hour at 4°C. The sample was then washed twice with 5CV of wash buffer (25 mM Hepes, 70 mM NH_4_Cl, 30 mM KCl, 7 mM MgCl_2_, 10% glycerol, 40 mM imidazole, 4 mM beta-mercaptoethanol) and protein was eluted with 5 times 1 CV of elution buffer (25 mM Hepes, 70 mM NH_4_Cl, 30 mM KCl, 7 mM MgCl_2_, 10% glycerol, 250 mM imidazole, 4 mM beta-mercaptoethanol). Purified protein was concentrated (Amicon® Ultra 10K MWCO) and rebuffered with storage buffer (25 mM Hepes, 70 mM NH_4_Cl, 30 mM KCl, 7 mM MgCl_2_, 10% glycerol) in 1 mL final volume. Then 1 mL 100% glycerol was added and protein was flash frozen and stored at −80°C.

#### Bpa photocrosslinking

To crosslink VDAC3 to FAM136A as shown in **Fig. 3A-C; Extended Data Fig. 5B**, VDAC3 constructs harboring amber stop codon mutations at positions I29 and N57 (VDAC3*) were transcribed as described above. The transcription product was then translated in the RRL system supplemented with ^35^S-methionine, Bpa-tRNA(UAG) (4 μM final concentration), Bpa (0.1 mM final concentration) and Bpa aminoacyl synthetase (5 μM final concentration). VDAC3* was translated in the absence or presence of 10 μM 3xFLAG-FAM136A. The translation product was run over a sucrose gradient as described above and each collected fraction was photo crosslinked via exposure to UV light for 10 min. The samples were denatured with 1% SDS and further analyzed via denaturing FLAG-IP as described above.

### In vitro reconstitution of VDAC3 insertion into the outer mitochondrial membrane (OM)

Isolation of mitochondria from K562 cells and in vitro insertion reactions shown in **Fig. 3D-F; Extended Data Fig. 6A and C** were conducted as described previously^83^.

#### Isolation of mitochondria from K562 cells

Mitochondria were isolated based on an adaption from an established protocol^87^ as described previously^30,83^. Briefly, 240 mL of K562 WT or FAM136A KO cells grown to 1×10^6^ cells/mL were centrifuged for 5 min at 300x g, and the pellet was washed once in homogenization buffer (210 mM mannitol, 70 mM sucrose, 5 mM HEPES-KOH pH 7.5, 1 mM EDTA, cOmplete^TM^ EDTA-free Protease Inhibitor Cocktail (Roche)). The weight of the resulting cell pellet was calculated, and 2 mL of homogenization buffer was added per 0.5 g of pellet. After resuspension, cells were incubated on ice for 10 min, before a 2 mL glass Dounce homogenizer with a tight-fitting pestle was used to lyse cells using ∼30 passes. The lysate was centrifuged at 1300x g for 5 min, and the supernatant was transferred to a clean tube to remove nuclei and unlysed cells. This step was repeated twice, before mitochondria were pelleted by centrifugating at 11,000x g for 10 min. Pelleted mitochondria were washed twice in isolation buffer (210 mM mannitol, 70 mM sucrose, 5 mM HEPES-KOH pH 7.5, 1 mM EDTA) and the final pellet was resuspended in 15-20 µL of isolation buffer. Prior to downstream use, total mitochondrial protein concentration was measured using a Bradford assay and normalized to 5 mg/mL for both WT and FAM136A KO mitochondria.

#### In vitro insertion into purified mitochondria

In vitro transcriptions and translations were performed in RRL as described above. Insertion reactions were performed post-translationally as described previously^30,83^. Mitochondria purified from either WT or FAM136A KO K562 cells were diluted with import buffer (20 mM HEPES-KOH, pH 7.5, 250 mM sucrose, 5 mM magnesium acetate, 20 mM potassium acetate, 2.5 mM ATP, 15 mM succinate) to 0.3 mg/mL and 0.08X (v/v) translation product was added. Insertion reactions were incubated at 32 °C for 5, 10, 20 or 30 min.

#### Mitochondrial swelling and PK digest

To differentiate between the population of nascent VDAC integrated into the OM vs the IMS, we isolated the IMS fraction after our in vitro translation and insertion reactions. To swell mitochondria after integration of the in vitro translated protein, mitochondria were first pelleted for 5 min at 11,000 × g at 4 °C to remove non-inserted translation product and then resuspended in 11x volume of swelling buffer (10 mM HEPES, pH7.2, 1 mM EDTA) and incubated for 25 min at 4 °C. While in this sample the outer membrane is ruptured, it contains the total amount of inserted translation product. We therefore call this sample “total”.

Using this swelling approach the membrane is not fully dissolved but fractured generating mitoplasts and outer membrane proteins that were successfully inserted into the outer membrane, should be protected from enzymatic digestion (**Extended Data Fig. 6A**)^43^. Protection from pyruvate kinase (PK) was assessed by adding 1 uL PK to 39 uL reaction and incubated on ice for one hour. The reaction was quenched by the addition of 0.8 µl 250 mM PMSF in DMSO and incubated on ice for 3 min. The whole reaction was then transferred to boiling 60 µl 1.67% SDS / 0.1 M Tris pH 8.0 and boiled for 3 min. Samples were prepared by adding sample buffer and analyzed via SDS-PAGE and autoradiography.

#### Quantification of insertion reaction

Gels were first analyzed using the ImageJ Gel analyzer measuring the intensity of each band. The intensity for 0 min was set to 0 and intensities were fit with a one-phase association fit (Eq. 3) for which Y starts at Y_0_, then goes up to Plateau with one phase.

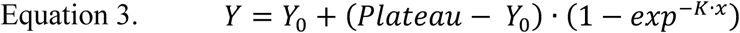

In which *Y*_0_ = 0, *K* is the rate constant in units that are the reciprocal of the x axis units and *x* is the time in minutes.

*Plateau* reflects the maximum amount of inserted substrate and the WT *Plateau* (*Plateau*_W*T*_) value was then used to normalize the data for comparison by calculating the fraction of inserted substrate at each time point under each condition by setting (*Plateau*_W*T*_) to be 1 and *Y*_0_ = 0.

### Sequence alignments

Sequence alignment of FAM136A and the TIMs in **Fig. 1D** was produced using MUSCLE^88^ and visualized using the ESPript 3.2 server^89^. Sequence alignment of VDAC1,2 and 3 and Porin1 and Porin 2 in **Extended Data Fig. 8A** was performed via the Jalview software using Clustal^90^ with default parameters and visualized using Jalview^91^.

### Structural analysis

Models of FAM136A monomer and dimer were generated using AlphaFold3^45^. Analysis and graphics that compared TIM9 and FAM136A structure in **Extended Data Fig. 2A** were performed using UCSF ChimeraX^92^. Structural representation of the hydrophobicity of each residue as shown in **Extended Data Fig. 7B** was performed by mapping the Gibbs free energy of unfolding of each amino acid determined by ^93^ onto the occupancy values in the pdb using tcl scripting in the VMD software^94^.Structural representation of the electrostatic potential as shown in **Extended Data Fig. 9B** was calculated using the APBS Electrostatics plugin from PyMOL^95^.Structural representation of the conservation of FAM136A as shown in **Fig. 6B** was calculated using the ConSurf webserver^66–68^ and visualized using PyMOL^95^.

### Hydrophobicity calculation

Hydrophobicity values were calculated by adding the Gibbs free energy of unfolding of each amino acid determined by^93^ and dividing by the total number of residues in the molecule.

### Evolutionary tree

To investigate if FAM136A is conserved across species, we queried the Interpro data base family (IPR008560) for FAM136A in different species across the phylogenetic tree. We find that FAM136A homologs exist in plants and metazoans as well as in choanoflagellata but not in fungi or filasterea. Accordingly, we investigated the amount of VDAC isoforms/paralogs in representative species for metazoans (vertebrates: H. sapiens; M. musculus; X. tropicalis; S. salar; and invertebrates: O. dentatum; D. melanogaster), Choanoflagellata (S. rosetta), Filasterea (C. owczarzaki) as well as fungi (S. cervisiae; N. crassa) and plants (A. thaliana; P. trichocarpa) and the total number of genes in the genome of each species.

## EXTENDED DATA FIGURES

**Extended Data Fig. 1:**
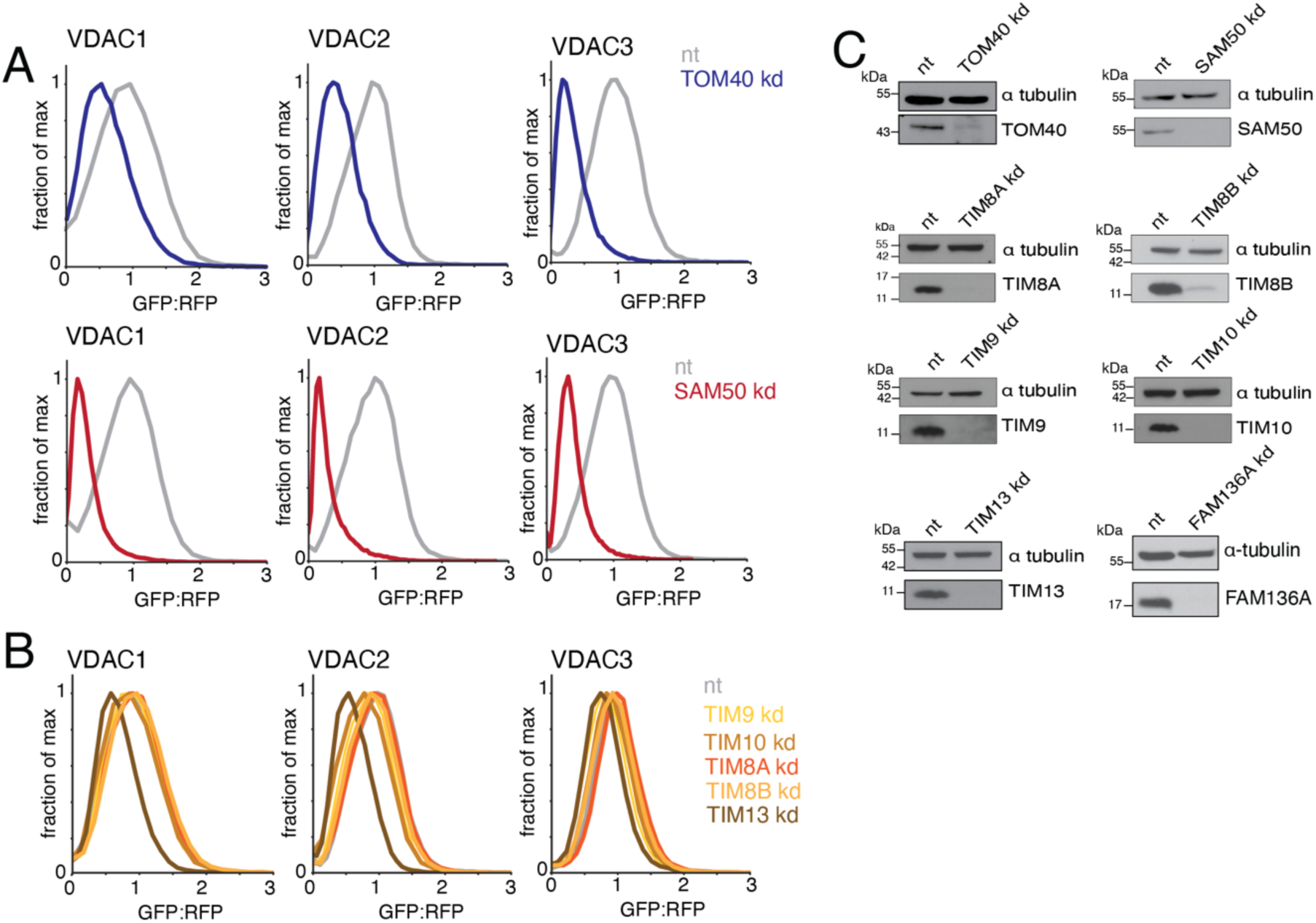
Validation of the split GFP system for studying the biogenesis of VDACs in human cells. **(A)** We tested if the split GFP reporter system described in Fig. 1A faithfully recapitulated known aspects of VDAC biogenesis. To do this, we used the GFP11-VDAC reporters for all three mammalian VDAC paralogs (VDAC1-3) and tested whether their integration into mitochondria depended on both the translocase of the outer membrane (TOM) complex and the sorting and assembly machinery (SAM). Integration into mitochondria of the GFP11-VDAC1-3 reporters was assessed in human K562 CRISPRi cells after depletion of the two central components of these complexes, TOM40 via shRNA, or SAM50 via sgRNA (kd) compared to a non-targeting control (nt). 48 h after transduction with either the shRNA or sgRNA, cells were treated with puromycin for three days to select for cells expressing the shRNA/sgRNA. After six days of depletion, cells were transduced with the indicated reporters and analyzed by flow cytometry on day eight. GFP fluorescence relative to a normalization marker (RFP) was determined using flow cytometry and the GFP:RFP signals are displayed as histograms, which are normalized such that GFP:RFP ratio of the nt control is set to 1. **(B)** As in (A) but to determine the dependence of the three GFP11-VDAC1-3 reporters on the known IMS-resident chaperones, TIM9/10 and TIM8/13. Mitochondrial integration of GFP11-VDAC1,2, and 3 was assessed in human K562 CRISPRi cells upon depletion of TIM8A, TIM8B, TIM9, TIM10, TIM13 (kd) compared to a non-targeting control (nt). Displayed are representative normalized histograms for the mitochondrial integration of each reporter (GFP), relative to an expression control (RFP). Quantification of this experiment is displayed in Fig. 1B. **(C)** Cells from (A), (B) and Fig. 1E-G were subjected to western blot analysis to assess successful depletion of TOM40, SAM50, the small TIMs, and FAM136A. Human K562 CRISPRi cells were transduced with either TOM40 shRNA, SAM50 sgRNA, TIM sgRNA, FAM136A sgRNA or a non-targeting control shRNA/sgRNA harboring puromycin and BFP selection markers. Following puromycin selection and eight days of depletion, cells were lysed with 1% GDN and the total protein fraction (20 μg/lane) were analyzed via SDS-PAGE and western blot. α-tubulin was used as loading control

**Extended Data Fig. 2:**
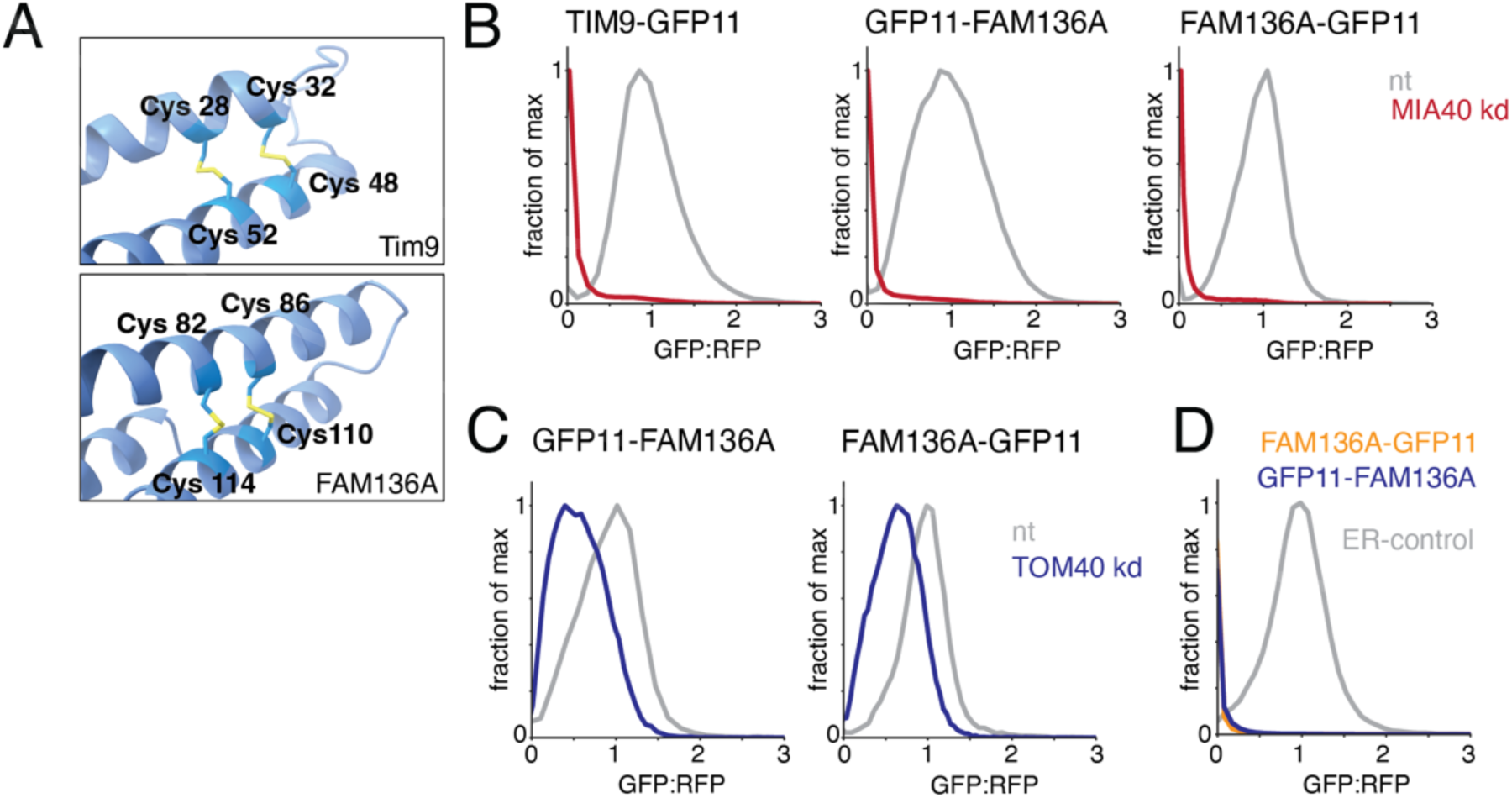
FAM136A localizes to the IMS and contains similar sequence features to the TIM chaperones. **(A)** Structural comparison between the CX_3_C motif of human TIM9 and FAM136A. Shown are excerpts of the experimental TIM9 (PBD# 2BSK) and AlphaFold3 predicted FAM136A (AF-Q96C01-F1-v6) structures in cartoon representation with the CX_3_C motif cysteines are highlighted as stick representation. **(B)** Given the presence of a twin CX_3_C motif, we hypothesized that FAM136A may depend on MIA40 for its folding in the IMS, as has been reported previously for the TIMs and other disulfide-dependent proteins^34,98^. To test this, we generated a GFP11-tagged FAM136A reporter, for use in the split GFP reporter system described in Fig. 1A. Integration into mitochondria of FAM136A tagged on either termini with GFP11 or the control TIM9-GFP11 was assessed in human K562 CRISPRi cells that constitutively expresses GFP1-10 in the IMS upon depletion of MIA40 via sgRNA (kd) compared to a non-targeting control (nt). GFP fluorescence relative to a normalization marker (RFP) was calculated and the GFP:RFP ratio was displayed as histograms normalized to the nt control. **(C)** As in (B) except for depletion of TOM40 via shRNA (kd) compared to a non-targeting control (nt). **(D)** The split GFP reporter system described in Fig. 1A with modifications was used to test if FAM136A mislocates to the ER. Here, integration into the ER of FAM136A tagged on either termini with GFP11 or the control OMP25-GFP11 reporter was assessed in human K562 CRISPRi cells that constitutively expresses GFP1-10 in the ER^76^. GFP fluorescence relative to a normalization marker (RFP) was calculated and the GFP:RFP signals are displayed as histograms

**Extended Data Fig. 3:**
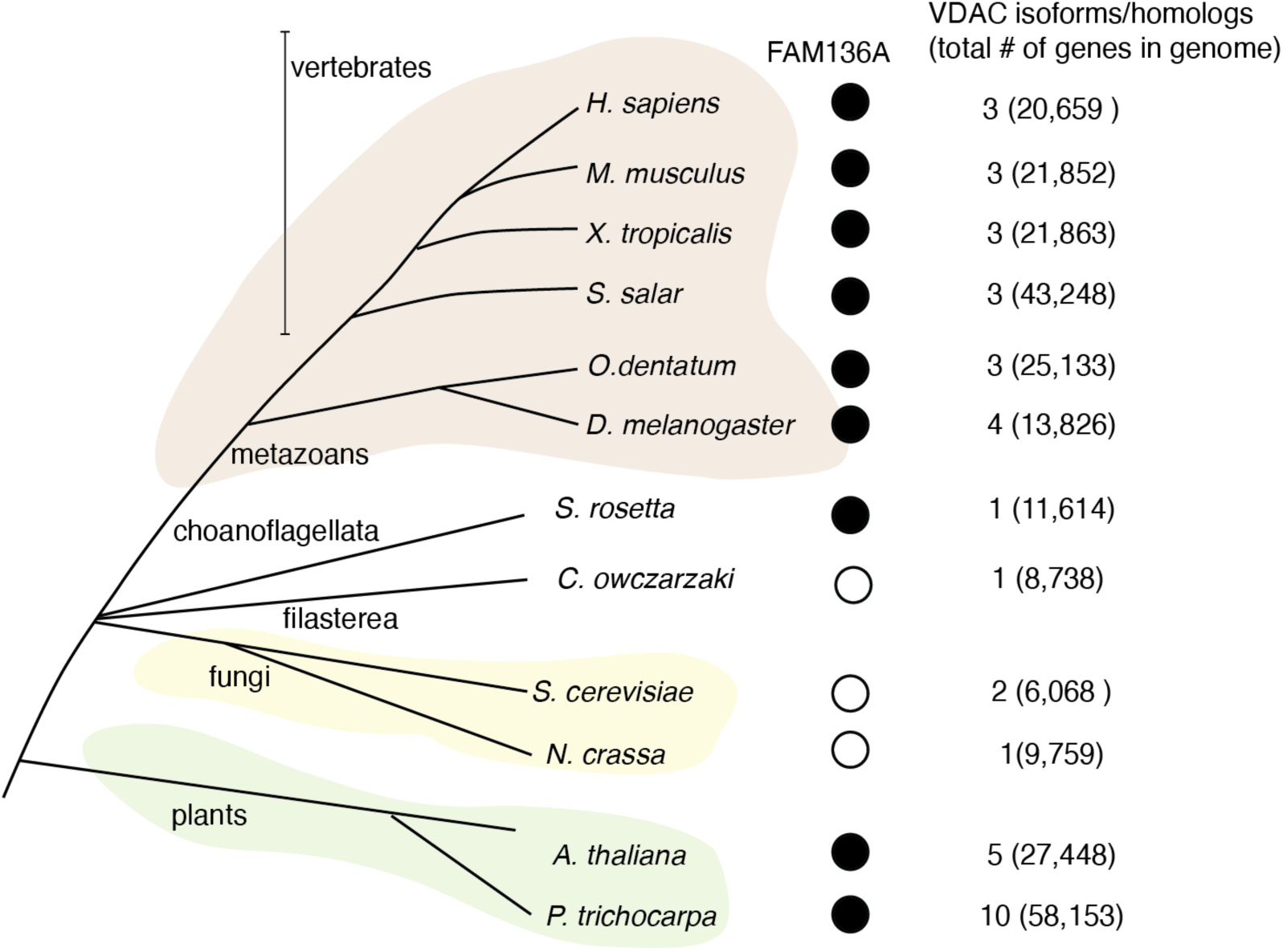
FAM136A is conserved in metazoans and plants. To investigate FAM136A conservation across species, we queried the Interpro data base family (IPR008560) for FAM136A. We only found FAM136A in eukaryotes and picked representative members of the Opisthokonta clade including Metazoans (both vertebrates and invertebrates) (vertebrates: *H. sapiens; M. musculus; X. tropicalis; S. salar; and invertebrates: O. dentatum; D. melanogaster*), Choanoflagellata (*S. rosetta*), Filasterea (*C. owczarzaki*) and Fungi (S*. cervisiae; N. crassa*) as well as representative members of the plants (*A. thaliana; P. trichocarpa*). For each organism, we searched for FAM136A in their proteomes (Uniprot). A filled dot indicates that FAM136A exists in the species’ proteome while an unfilled dot indicates that no FAM136A homolog was found in the Uniprot data base. Also displayed are the number of VDAC paralogs in each representative species along with the total number of genes in the genome. While FAM136A is broadly conserved across metazoan, and plants and select choanoflagellates, it has been lost in the fungal lineage. This pattern of conservation could be consistent with FAM136A being present in the latest eukaryotic common ancestor (LECA), but does not seem to have been evolutionarily required in fungus, which have a small mitochondrial genome and express fewer VDAC paralogs^2,4^.

**Extended Data Fig. 4:**
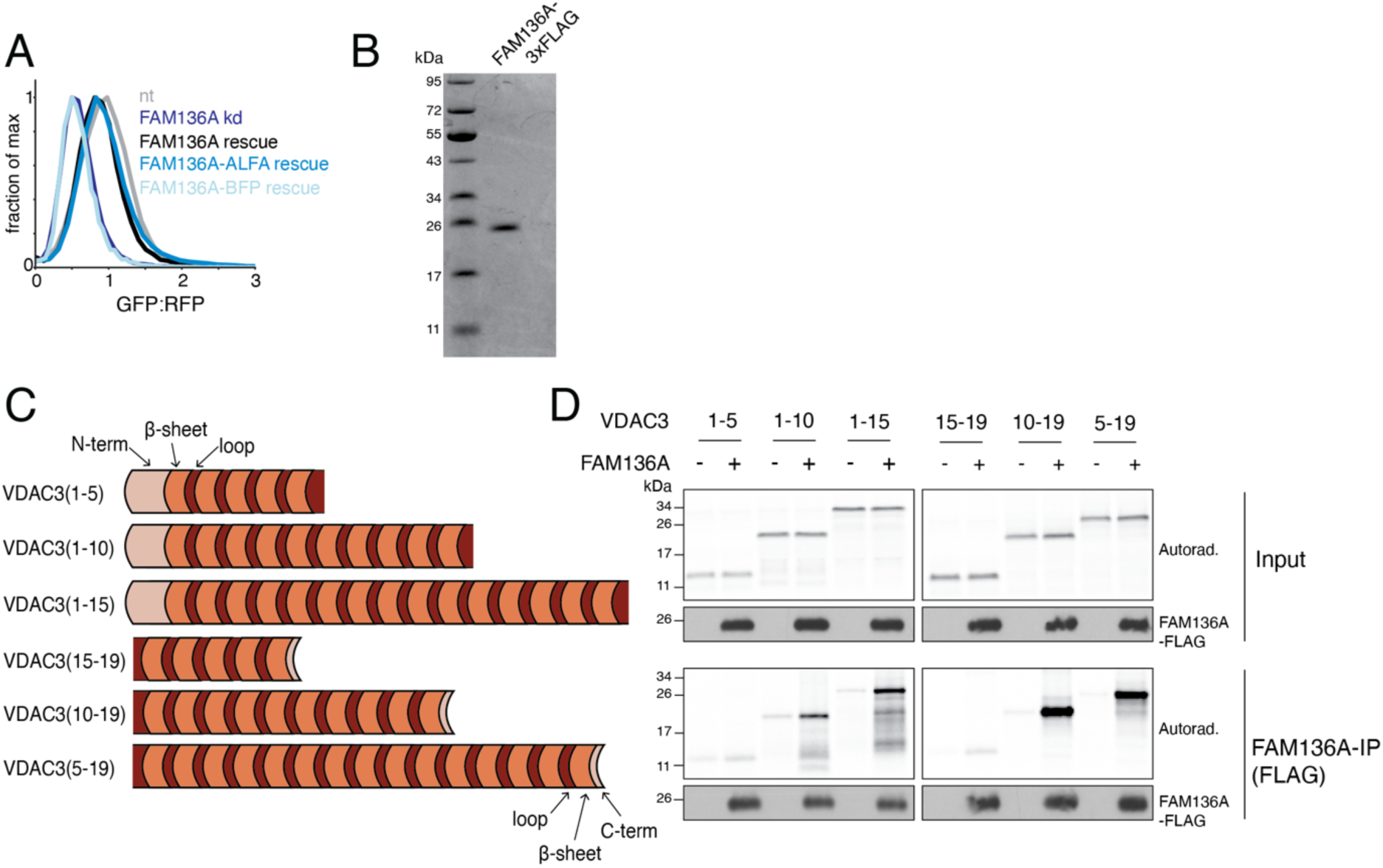
Molecular analysis of FAM136A in VDAC3 biogenesis. **(A)** To determine which affinity tags do not affect the function of FAM136A, we leveraged our ratiometric fluorescent VDAC3 reporter. Mitochondrial integration of GFP11-VDAC3 was assessed in human K562 CRISPRi cells that constitutively expresses GFP1-10 in the IMS upon depletion of FAM136A (kd) compared to a non-targeting control (nt). Alongside the reporter, FAM136A rescue constructs with either no tag, an ALFA tag, or fused to a BFP were transduced. GFP fluorescence relative to a normalization marker (RFP) was calculated and the GFP:RFP signals are displayed as histograms. Only the untagged or ALFA-tagged FAM136A was able to rescue the kd phenotype. We hypothesized that fusion with a full length BFP may sterically affect the ability of FAM136A to translocate into the IMS and/or perform its function. Further experiments that required FAM136A tagging in cells therefore relied on this ALFA-tagging strategy, while in vitro experiments utilized a 3xFLAG tag in the same position. **(B)** In order to perform the in vitro assays (Fig. 2D; Fig. 3B-C; Fig. 4B,D,E; Fig. 5B,D; Extended Data Fig. 4D; Extended Data Fig. 5B-D; Extended Data Fig. 7A,D,F; Extended Data Fig. 9A; Extended Data Fig. 10D), recombinant FAM136A-3xFLAG was expressed and purified from *E. coli*. Briefly, FAM136A-3xFLAG was expressed with a N-terminal His14-SUMO^Eu1^ tag and was captured by Ni-NTA agarose after the cells were lysed by sonication. Proteins were eluted by SUMO^Eu1^ protease cleavage and further purified using a Superdex 200 increase 3.2/300 size exclusion column. Shown is the final protein sample analyzed on SDS-PAGE and stained with Coomassie Blue. **(C)** Schematic of the VDAC3 truncation mutants used in (D). To test for the minimum length of VDAC3 needed for binding to FAM136A, we generated a series of successive truncation mutants of VDAC3. VDAC3 was either truncated from the C-terminus after the indicated b-strands or truncated from the N-terminus before the indicated b-strands. Note in all cases, the intervening loops between each b-strand remained intact in these constructs. VDAC3 is shown in orange, with b-strands and intervening soluble loops displayed in light and dark shades, respectively. **(D)** Analysis of FAM136A binding of the nascent VDAC3 truncation mutants shown in (C) in vitro. The ^35^S-methionine labeled substrates were translated in rabbit reticulocyte lysate (RRL) in the absence or presence of recombinant FAM136A-3xFLAG purified from *E. coli*. FAM136A was immunoprecipitated using anti-FLAG resin and eluted with 3xFLAG peptide. Co-purification of each substrate was analyzed by SDS-PAGE and autoradiography. Samples were also subjected to western blotting to ensure an equal amount of FAM136A-3xFLAG in the input and elution. We found that as few as five b-strands and their intervening loops were sufficient for binding to FAM136A, though ten or more were captured more efficiently. Because all truncations across VDAC3 were able to bind FAM136A, we concluded that there is not a single specific binding site of FAM136A for VDAC3.

**Extended Data Fig. 5:**
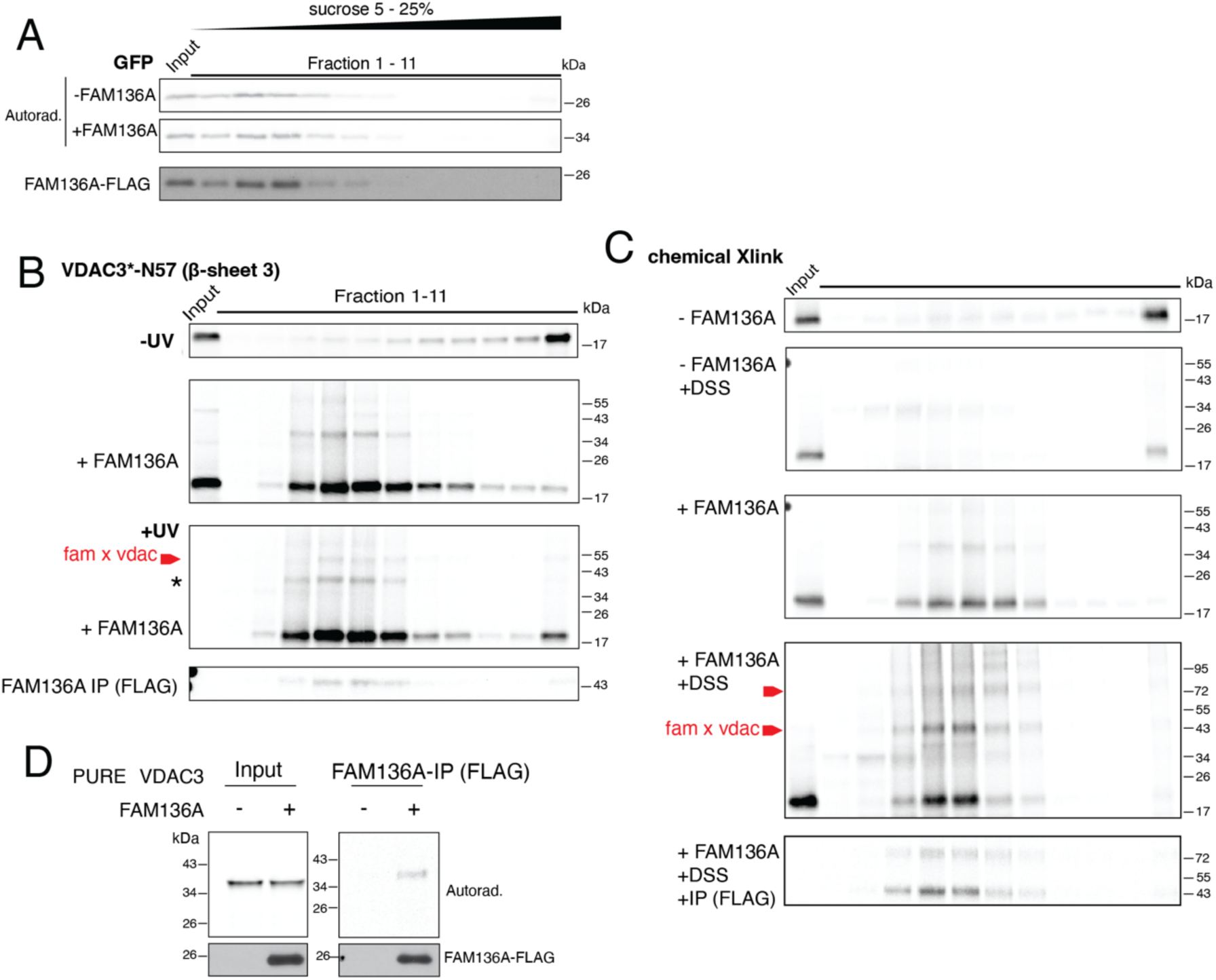
FAM136A solubilizes VDAC3 via a direct interaction. **(A)** Analysis of GFP in the absence or presence of FAM136A-3xFLAG as a control for Fig. 3B. ^35^S-methionine labeled GFP was translated in RRL in the absence or presence of recombinant FAM136A-3xFLAG purified from *E. coli* and fractionated on a sucrose gradient as described in **Fig. 3A**^65^. Nascent GFP was visualized by SDS-PAGE and autoradiography, while the migration of FAM136A-3xFLAG on the sucrose gradient was determined by western blotting. Unlike VDAC3, GFP migration is not affected by the presence of FAM136A, consistent with earlier experiments showing that GFP does not bind FAM136A (Fig. 2D). GFP, a soluble β-barrel migrates at a fraction consistent with its 40 kDa size. **(B)** Site-specific UV crosslinking of VDAC3* and FAM136A. To test another site for site-specific crosslinking along placing the crosslinker into the first β-sheet, VDAC3 construct harboring an amber stop codon mutation at position N57 (in the third β-sheet) (VDAC3*) (1-10) was translated in the presence of absence of FAM136A-3xFLAG. Translations were then fractionated on a sucrose gradient (Fig. 3B). Each fraction was then subjected to photo crosslinking under UV light, followed by a FLAG-IP under denaturing conditions to enrich for FAM136A crosslinked species. Samples were analyzed by SDS-PAGE and autoradiography. **(C)** Chemical crosslinking of VDAC3 with FAM136A shows multiple crosslinked bands suggesting multiple binding sites. ^35^S-methionine labeled VDAC3(1-10) translated in rabbit reticulocyte lysate (RRL) in the absence of presence of FAM136A-3xFLAG and fractionated via a sucrose gradient as described in Fig. 3A. Each fraction was incubated with DSS, a primary amine specific crosslinker, to test for interactions between VDAC3 and FAM136A-3xFLAG. Finally FLAG-IP were conducted on each sample to enrich the crosslinked product. All samples were analyzed by SDS-PAGE and autoradiography. Shown are samples in the absence and presence of FAM136A as well as the crosslinker. In the IPed sample, there are two bands corresponding to the molecular weight of one and two FAM136A-3xFLAG bound to VDAC3(1-10). **(D)** FAM136A purified from *E.coli* binds VDAC3 in vitro without the need for additional factors. To test if the FAM136A-VDAC3 interaction required other factors for substrate loading of VDAC3 onto FAM136A, we leveraged the PURE system, which contains only the purified recombinant *E.coli* translation machinery, but no chaperones or eukaryotic factors^80^. ^35^S-methionine labeled VDAC3 was translated in the PURExpress system in the absence or presence of recombinant FAM136A-3xFLAG purified from *E. coli*. FAM136A was immunoprecipitated using anti-FLAG resin and eluted with 3xFLAG peptide. Co-purification of VDAC3 was analyzed by SDS-PAGE and autoradiography. Samples were also subjected to western blotting to ensure an equal amount of FAM136A-3xFLAG in the input and elution.

**Extended Data Fig. 6:**
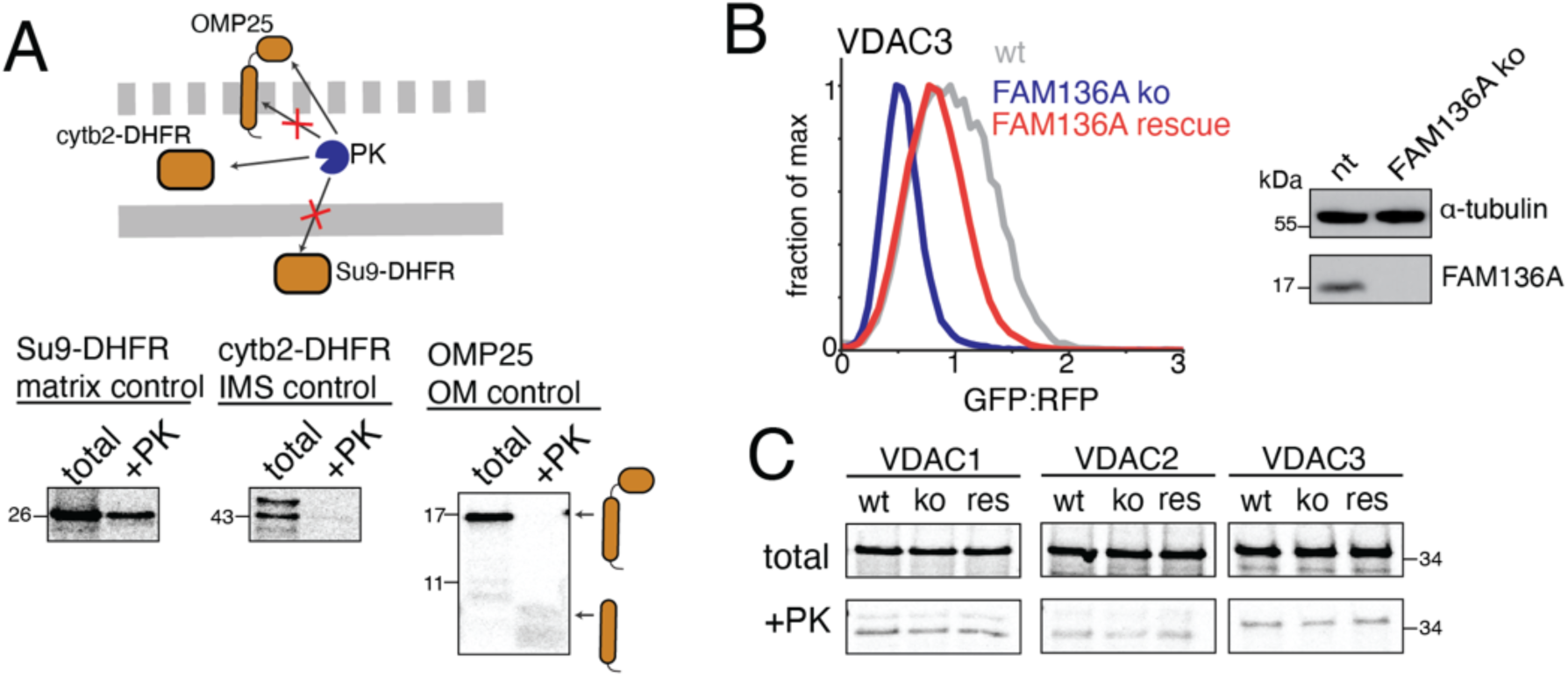
FAM136A is required for OM insertion of the VDACs. **(A)** To identify which step in VDAC biogenesis is mediated by FAM136A, we leveraged an in vitro insertion assay. Shown here are matrix, IMS and OM controls validating the experiment described in Fig. 3D. In each case we used untagged proteins to exclude potential tagging artifacts. The indicated substrates were translated in rabbit reticulocyte lysate (RRL) in the presence of ^35^S-labeled methionine and then incubated with mitochondria purified from human K562 CRISPRi cells. Mitochondria were separated from any non-integrated translation products by centrifugation. To differentiate between substrate in the IMS and the OM, mitochondria were swollen using hypotonic buffer to permeabilize the OM without solubilizing, thereby creating mitoplasts. In these mitoplasts, the IMS becomes accessible, while the inner membrane remains intact. They were then treated with proteinase K (PK) to digest non-protected fragments. Under these conditions, (1) nascent substrates within the IMS become protease accessible, (2) nascent substrates in the matrix remain protease inaccessible and (3) those integrated into the OM are protected along their TM domain while cytosolic or IMS exposed domains become protease accessible. Shown are three control substrates before and after PK digest analyzed via SDS-PAGE and autoradiography. The matrix control Su9-DHFR is protected, no change in the fragment is observed. The IMS control cytb2-DHFR shows a loss of signal after PK treatment while the OM control OMP25 shows a shift in the observed band according to the cytosolic domain being digested while the TM domain remains protease protected. **(B)** To verify that the effects of FAM136A depletion on the OM insertion of VDAC in vitro (Fig. 3F) were specific we tested if we could rescue this phenotype by overexpression of exogenous FAM136A. Using our ratiometric fluorescent reporter system, we measured the mitochondrial integration of GFP11-VDAC3 in human K562 CRISPRi cells in which FAM136A has been knocked out (ko) compared to wild type cells (wt). Alongside the reporter, a FAM136A rescue construct was transduced along with a BFP to identify transduced cells. GFP fluorescence relative to a normalization marker (RFP) was calculated and the GFP:RFP signals are displayed as histograms. (Right) Wt and FAM136A ko cells were subjected to western blot analysis to assess successful depletion of FAM136A. Cells were lysed with 1% GDN and the total protein fraction (20 μg/lane) were analyzed via SDS-PAGE and western blot. α-tubulin was used as loading control. **(C)** Autoradiography of ^35^S-methionine labelled VDAC1, VDAC2 or VDAC3 inserted into purified mitochondria from K562 WT cells, FAM136A KO cells or FAM136A KO cells overexpressing the FAM136A rescue construct exogenously. To generate this cell line, FAM136A KO cells were transduced with FAM136A_ALFA_P2A_BFP rescue construct and sorted for BFP+ cells to create a pure cell population that contains the rescue construct. Shown are samples of the total amount of VDAC1-3 inserted into mitochondria and the amount of PK digest protected VDAC1-3.

**Extended Data Fig. 7:**
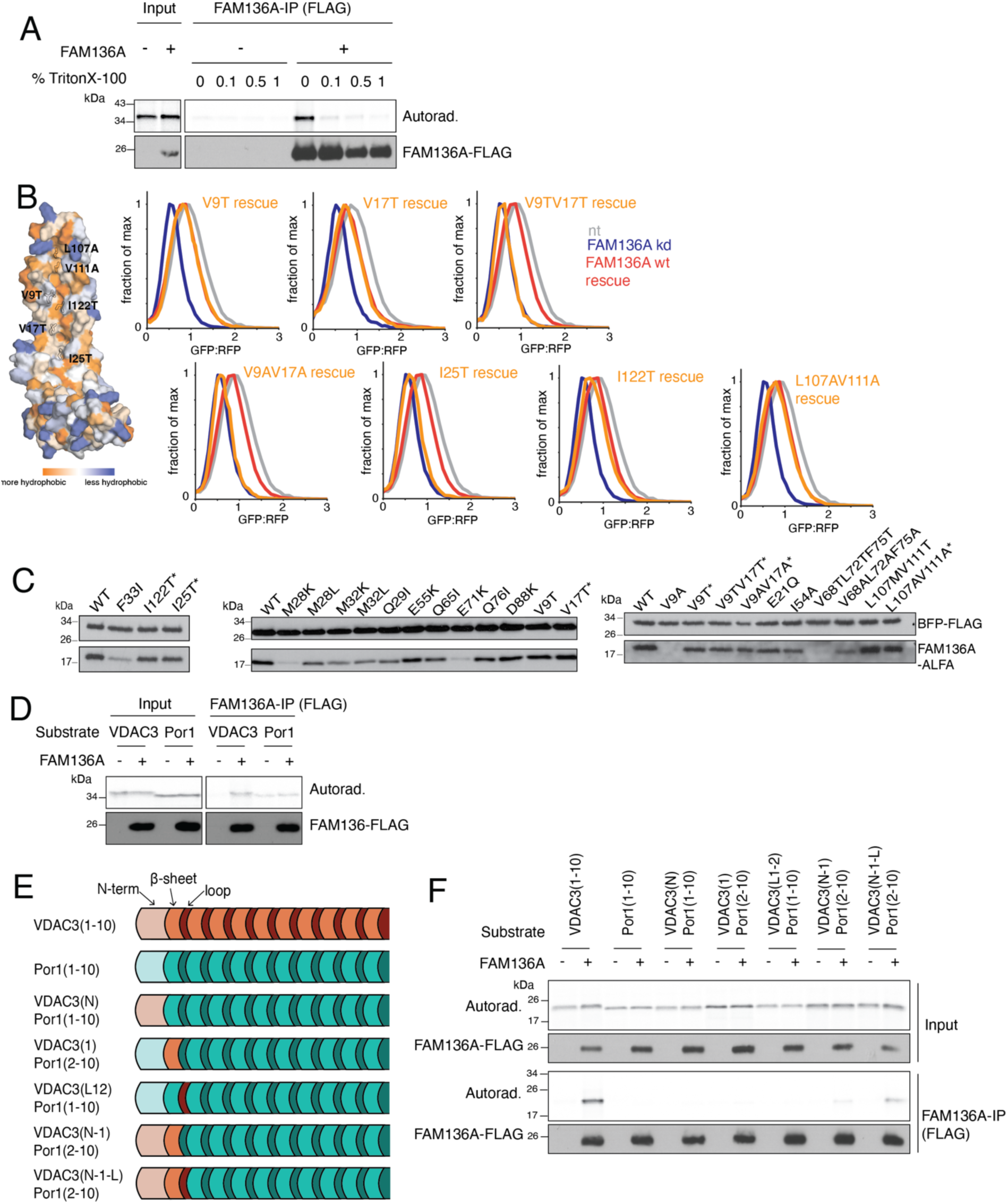
VDAC3 binds to the hydrophobic groove of FAM136A. **(A)** To determine the properties of the VDAC3-FAM136A interaction, we tested whether it was detergent sensitive in vitro. ^35^S-methionine labeled VDAC3 was translated in rabbit reticulocyte lysate (RRL) in the absence or presence of recombinant FAM136A-3xFLAG purified from *E. coli*. FAM136A was immunoprecipitated using anti-FLAG resin and eluted with 3xFLAG peptide after washing with increasing concentrations of TritonX-100. Co-purification of VDAC3 in each condition was analyzed by SDS-PAGE and autoradiography. Samples were also subjected to western blotting to ensure an equal amount of FAM136A-3xFLAG being immunoprecipitated. Loss of binding of VDAC3 to FAM136A at increasing concentrations of detergent is consistent with a hydrophobic interaction between FAM136A and VDAC3. **(B)** Mutational analysis of the role of the hydrophobic groove in FAM136A on VDAC3 biogenesis performed using the ratiometric fluorescent reporter system described in Fig. 1A. Wildtype (wt) FAM136A or the indicated mutants were transduced into FAM136A depleted K562 CRISPRi cells expressing GFP11-VDAC3. (Left) A space-filling representation of the predicted AlphaFold3 model of a FAM136A monomer (alphafoldid: AF-Q96C01-F1-v6) in which the indicated residues were mutated in silico using the CHARMM-GUI protein builder webtool^99^ is displayed with the hydrophobic residues colored according to their hydrophobicity value based on^93^. (Right) Integration into mitochondria of GFP11-VDAC3 was assessed in human K562 CRISPRi cells in which FAM136A has been depleted (kd) compared to a non-targeting control (nt). Alongside the reporter, a FAM136A mutant rescue construct tagged with ALFA_P2A_BFP was introduced into the cells. GFP fluorescence relative to a normalization marker (RFP) was calculated and the GFP:RFP signals are displayed as histograms. **(C)** Cells from (**B**) were subjected to western blot analysis to assess expression levels of the mutant rescue constructs compared to wt FAM136A. Human K562 CRISPRi cells were transduced with FAM136A sgRNA or a non-targeting (nt) control sgRNA harboring a puromycin selection marker. After two days, cells were treated with puromycin for three days to select for cells with integrated sgRNA. After five days cells were transduced with the GFP11-VDAC3 reporter alongside an ALFA-tagged FAM136A wt or mutant (expressed as an ALFA-FAM136A_p2a_BFP) or a BFP only control. BFP expression can then be used to identify cells transduced with the FAM136A rescue constructs. After eight days of depletion cells were lysed with 1% GDN. Samples were normalized first to the total amount of protein based on the A280 and then to the match the percentage of BFP to correct for varying transduction efficiencies. Samples were then analyzed via SDS-PAGE and western blot, for BFP-FLAG and FAM136A-ALFA. Based on these experiments we were able to identify FAM136A mutants where loss of function could not be explained by changes to FAM136A stability and/or expression and they are indicated with asterisk (*). **(D)** To test if Por1, the yeast homolog of the VDACs, is able to bind to FAM136A we leveraged our in vitro translation and immunoprecipitation assay as in Fig. 2D. The ^35^S-methionine labeled VDAC3 or Por1 were translated in RRL in the absence or presence of recombinant FAM136A-3xFLAG purified from *E. coli*. FAM136A was immunoprecipitated using anti-FLAG resin and eluted with 3xFLAG peptide. Co-purification of VDAC3 or Por1 was analyzed by SDS-PAGE and autoradiography. Samples were also subjected to western blotting to ensure an equal amount of FAM136A-3xFLAG in the input and elution. **(E)** To define the sequence requirements for capture by FAM136A we leveraged the observation that while truncated VDAC3 binds FAM136A, similar length Por1 cannot. In Fig. 4C, we found that fusion of the N-terminus of VDAC3 (including the soluble N-termini, first b-strand, and first intervening loop) was sufficient to confer binding of Por1. To further dissect the relative contributions of these features to FAM136A binding, we systematically fused each of these elements to our Por1(1-10) sequence. Shown is a schematic of resulting VDAC3-Porin1(1-10) chimera constructs used in (F). VDAC3 is shown in orange and Por1 in green, with their b-strands and intervening soluble loops displayed in light and dark shades, respectively. **(F)** ^35^S-methionine labeled substrates were translated in RRL in the absence or presence of recombinant FAM136A-3xFLAG purified from *E. coli*. FAM136A was immunoprecipitated using anti-FLAG resin and eluted with 3xFLAG peptide. Co-purification of each substrate was analyzed by SDS-PAGE and autoradiography. Samples were also subjected to western blotting to ensure an equal amount of FAM136A in the input and elution. We see the most pronounced binding for fusions that include the soluble N-terminus, first b-strand, and first loop of VDAC3. However, no single one of these elements alone is sufficient to confer binding.

**Extended Data Fig. 8:**
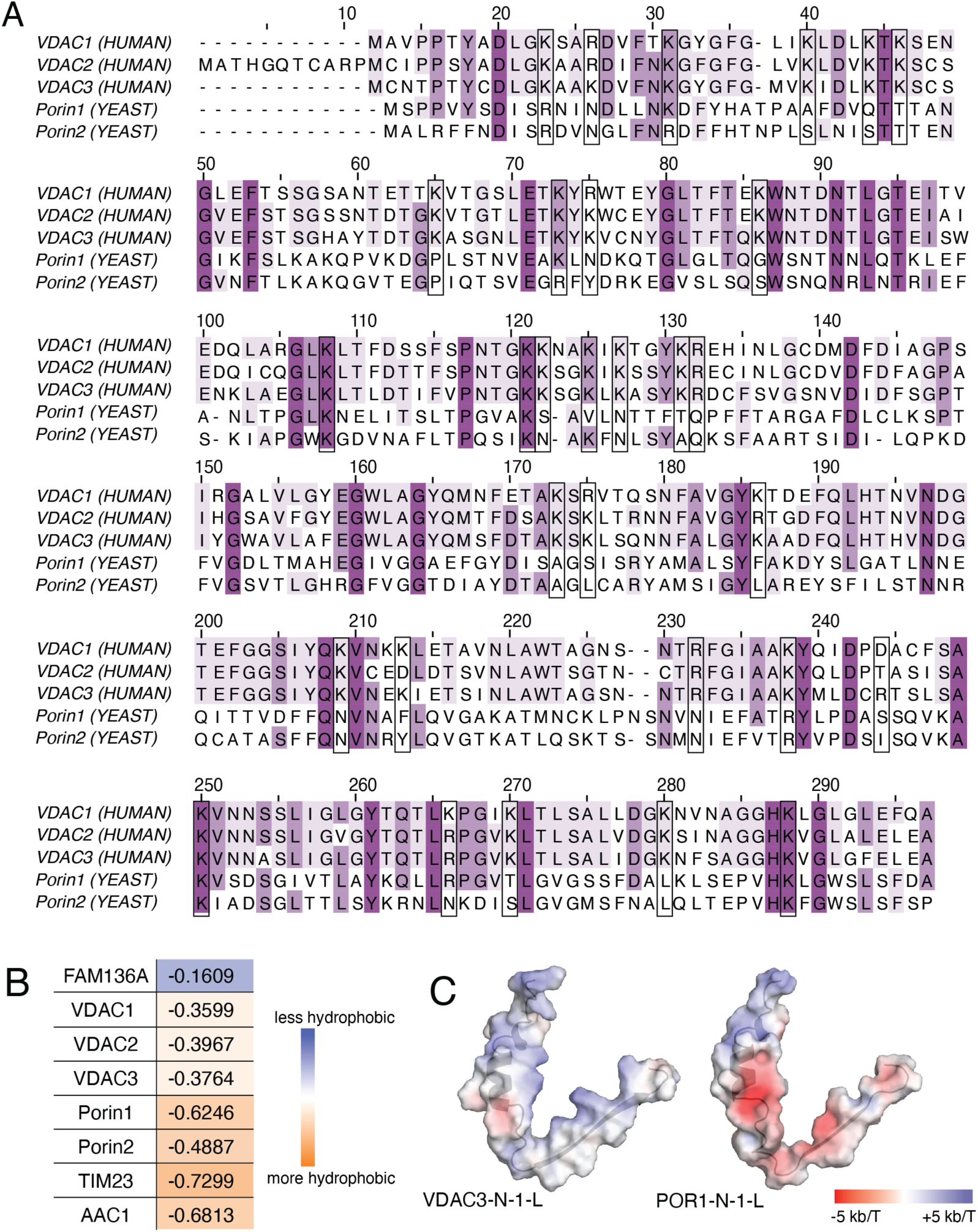
The VDACs and Porins contain distinct levels of charge. **(A)** Sequence alignment of human VDAC1-3 and yeast Porin 1 and 2. The level of conservation is shown in purple with darker colored residues being most conserved. Boxed are all positively charged lysine or argenine residues in VDAC3 and the corresponding residues in the paralogs. **(B)** Overall hydrophobicity of FAM136A, Porin1-2, VDAC1-3, AAC1 (IM Solute carrier 25; SLC25, a canonical TIM9/10 substrate), and TIM23 (the canonical TIM8/13 substrate). The sum of side chain transfer free energy for every amino acid was calculated and averaged over the number of residues of each protein^93^. Values are colored with orange representing higher overall hydrophobicity and blue representing overall lower hydrophobicity. **(C)** Comparison of the electrostatic potential of the N-termini of the minimal FMA136A binding site in VDAC3 with the corresponding sequence from Por1. A space-filling representation of the soluble N-terminus, first β-sheet, and first loop of the predicted AlphaFold3^45^ model of a FAM136A and the experimental structure of Por1 (PDB# 9JVQ) is displayed in which the residues are colored based on their electrostatic potential calculated via the APBS Electrostatics plugin for Pymol^95^. Por1 is enriched for negative charges in this region compared to VDAC3.

**Extended Data Fig. 9:**
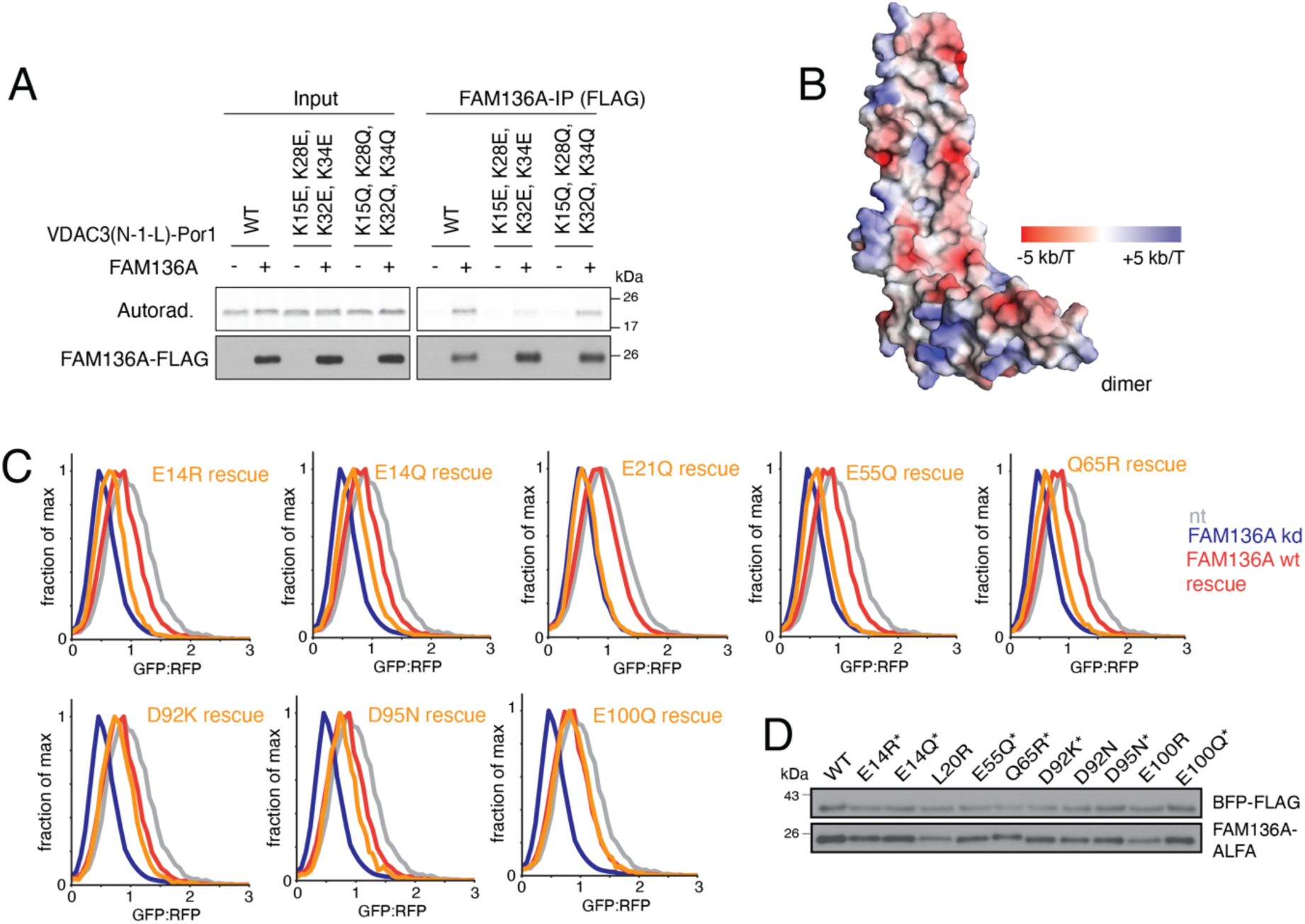
Substrate selectivity by FAM136A relies on the conserved negative charges that line its hydrophobic groove. **(A)** As in Fig. 4E, except to distinguish between whether VDAC3 binds FAM136A due primarily due to electrostatic attraction or whether Por1 is instead rejected from binding via electrostatic repulsion. Positively charged lysine residues in VDAC3 were replaced with either the negatively charged glutamic acid or neutral glutamine in the VDAC3(N-1-L)-Porin1(2-10) chimera (which contains a single FAM136A binding site). The ^35^S-methionine labeled constructs were translated in RRL in the absence or presence of recombinant FAM136A-3xFLAG purified from *E. coli*. FAM136A was immunoprecipitated under native conditions using anti-FLAG resin and eluted with 3xFLAG peptide. Co-purification of each mutant was analyzed by SDS-PAGE and autoradiography. Samples were also subjected to western blotting to ensure an equal amount of FAM136A-3xFLAG in the input and elution. While introduction of negative charge within the FAM136A binding site of the VDAC3(N-1-L)-Porin1(2-10) chimera markedly decreased binding, loss of positive charge also modestly affected binding. These data therefore suggest that both electrostatic repulsion and attraction may contribute to substrate selection by FAM136A. **(B)** Negative charges are enriched along the hydrophobic groove of FAM136A. A space-filling representation of the predicted AlphaFold3 model of the FAM136A monomer is displayed in which the residues are colored based on their electrostatic potential calculated via the APBS Electrostatics plugin for Pymol^95^. **(C)** To determine the role of the negatively charged residues in FAM136A, we performed mutational analysis using the ratiometric fluorescent reporter for VDAC3 (Fig. 1A). Shown here are representative data for the experiment summarized in Fig. 4F. Integration into mitochondria of GFP11-VDAC3 was assessed in human K562 CRISPRi cells in which FAM136A has been depleted (kd) compared to a non-targeting control (nt). Alongside the reporter, a FAM136A mutant rescue construct tagged with ALFA_P2A_BFP was introduced into the cells. GFP fluorescence relative to a normalization marker (RFP) was calculated for BFP positive cells and the GFP:RFP signals are displayed as histograms. **(D)** Cells from (C) were subjected to western blot analysis to assess expression levels of the mutant rescue constructs compared to wt FAM136A. Human K562 CRISPRi cells were transduced with FAM136A sgRNA or a non-targeting control sgRNA harboring a puromycin selection marker. After two days, cells were treated with puromycin for three days to select for cells with integrated sgRNA. After five days cells were transduced with the GFP11-VDAC3 reporter alongside a FAM136A mutant reporter tagged with ALFA_P2A_BFP. This results in an ALFA-tagged protein while BFP serves as an expression control. After eight days of depletion cells were lysed with 1% GDN. Samples were normalized first to the total amount of protein based on the A280 and then to BFP percentage to correct for transduction efficiency. Samples were then analyzed via SDS-PAGE and western blot, blotting with antibodies for BFP-FLAG and FAM136A-ALFA. Based on these experiments we were able to identify FAM136A mutants where loss of function could not be explained by changes to FAM136A stability and/or expression and they are indicated with asterisk (*). Note that the western blot for E21Q is shown in Extended Data Fig. 7C.

**Extended Data Fig. 10:**
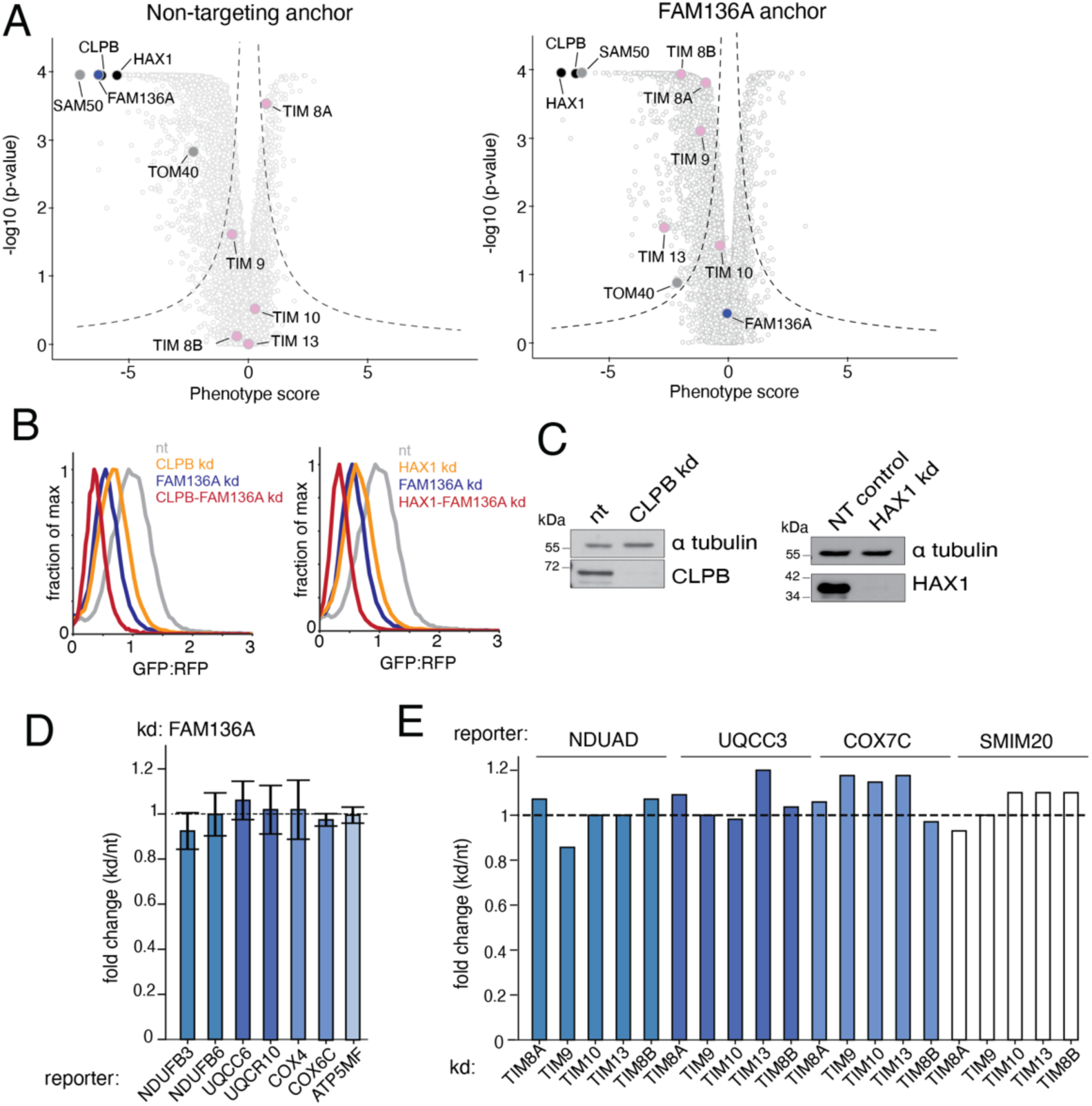
FAM136A works in parallel with other IMS chaperones and the ClpB disaggregase in VDAC3 biogenesis. **(A)** To understand how FAM136A fit in the larger context of the IMS chaperone network a genetic modifier screen to identify factors that have enhanced or diminished phenotypes on VDAC3 biogenesis in human cells upon depletion of FAM136A was performed using a genome wide CRISPRi dual-screen using the VDAC3 reporter depicted in **Fig. 1A**^46^. K562 CRISPRi cells stably expressing GFP11-VDAC3 along with the normalization marker (RFP) were transduced with a library harboring either a non-targeting control or FAM136A sgRNA as the genetic anchor. Following eight days of depletion, cells with altered GFP:RFP ratios were isolated using FACS and deep sequenced to identify enriched sgRNAs in the top (increased GFP:RFP) and bottom (decreased GFP:RFP) populations. Depicted are volcano plots of the phenotype score [log2(increased GFP:RFP/decreased GFP:RFP)] for the three strongest sgRNAs versus the -log of the Mann-Whitney p values for both the nt-anchored and FAM136A-anchored screens. Note the data from the nt-anchored screen has also been used to generate **Fig. 1C**, yet there, different genes are highlighted. Genes that fall outside the dashed lines represent statistically significant hits. Individual genes are displayed in grey, the biogenesis machineries TOM40 and SAM50 are highlighted in dark grey, the IMS-resident TIM chaperones in pink, CLPB and HAX1 are shown in black and FAM136A is shown in blue. Enhanced phenotypes in the FAM136A-anchored screen suggest synthetic effects, which would be indicative of factors in a parallel pathway. In contrast, diminished phenotypes in the FAM136A-anchored screen would suggest factors in the same pathway ^46^. **(B)** To study the genetic interaction between CLPB/HAX1 and FAM136A we performed genetic modifier experiments using the split GFP reporter system described in Fig. 1A. To do this we tested the effect of depletion of either HAX1 or CLPB alone or in combination with FAM136A on GFP11-VDAC3 biogenesis. In this assay, factors that function in parallel pathways would be expected to have an additive effect, while factors that function together in the same pathway would have the same phenotype as the single depletion alone. Integration into mitochondria of GFP11-VDAC3 was assessed in human K562 CRISPRi cells that constitutively expresses GFP1-10 in the IMS upon depletion of FAM136, CLPB, HAX1 single or FAM136A-CLPB or FMA136A-HAX1 dual knockdown sgRNA (kd) compared to a non-targeting control (nt). GFP fluorescence relative to a normalization marker (RFP) was calculated and the GFP:RFP signals are displayed as histograms. The additive effects of CLPB/HAX1 depletion with FAM136A would be most consistent with these factors functioning in parallel, partially redundant pathways. **(C)** Cells from (B) were subjected to western blot analysis to assess successful depletion of CLPB and HAX1. Human K562 CRISPRi cells were transduced with either CLPB or HAX1 sgRNA or a non-targeting control sgRNA harboring puromycin and BFP selection markers. After two days, cells were treated with puromycin for three days to select for cells with integrated sgRNA. After eight days of depletion cells were lysed with 1% GDN and the total protein fraction (20 μg/lane) were analyzed via SDS-PAGE and western blot, blotting with antibodies for α-tubulin as control as well as anti-CLPB and anti-HAX1 antibodies. **(D)** Not all inner mitochondria membrane located single TM reporters are affect by FAM136A depletion. The reporter system described in Fig. 1A was used to test the dependence of a wider range of inner mitochondria membrane located single TM reporters on depletion of FAM136A. Mitochondrial integration of GFP11-tagged reported substrates was assessed in human K562 CRISPRi cells upon depletion of FAM136A (kd) compared to a non-targeting control (nt). Shown is the fold change for each reporter calculated as (GFP:RFP in kd cells)/(GFP:RFP in nt cells). A fold change of 1 indicates no change compared to the nt control. Shown is the mean of 2-3 independent biological replicates ± SD. **(E)** Inner mitochondria membrane located single TM reporters that were FAM136A dependent (Fig. 5C) are not affect by small TIMs depletion. The reporter system described in Fig. 1A was used to test the dependence of a wider range of inner mitochondria membrane located single TM reporters on depletion of the small TIMs. Mitochondrial integration of GFP11-tagged reported substrates was assessed in human K562 CRISPRi cells upon depletion of TIM8A, TIM8B, TIM9, TIM10, TIM13 (kd) compared to a non-targeting control (nt). Shown is the fold change for each reporter calculated as (GFP:RFP in kd cells)/(GFP:RFP in nt cells). A fold change of 1 indicates no change compared to the nt control.

